# A Comprehensive Phylogenomic Platform for Exploring the Angiosperm Tree of Life

**DOI:** 10.1101/2021.02.22.431589

**Authors:** William J. Baker, Paul Bailey, Vanessa Barber, Abigail Barker, Sidonie Bellot, David Bishop, Laura R. Botigué, Grace Brewer, Tom Carruthers, James J. Clarkson, Jeffrey Cook, Robyn S. Cowan, Steven Dodsworth, Niroshini Epitawalage, Elaine Françoso, Berta Gallego, Matthew G. Johnson, Jan T. Kim, Kevin Leempoel, Olivier Maurin, Catherine McGinnie, Lisa Pokorny, Shyamali Roy, Malcolm Stone, Eduardo Toledo, Norman J. Wickett, Alexandre R. Zuntini, Wolf L. Eiserhardt, Paul J. Kersey, Ilia J. Leitch, Félix Forest

## Abstract

The tree of life is the fundamental biological roadmap for navigating the evolution and properties of life on Earth, and yet remains largely unknown. Even angiosperms (flowering plants) are fraught with data gaps, despite their critical role in sustaining terrestrial life. Today, high-throughput sequencing promises to significantly deepen our understanding of evolutionary relationships. Here, we describe a comprehensive phylogenomic platform for exploring the angiosperm tree of life, comprising a set of open tools and data based on the 353 nuclear genes targeted by the universal Angiosperms353 sequence capture probes. This paper (i) documents our methods, (ii) describes our first data release and (iii) presents a novel open data portal, the Kew Tree of Life Explorer (https://treeoflife.kew.org). We aim to generate novel target sequence capture data for all genera of flowering plants, exploiting natural history collections such as herbarium specimens, and augment it with mined public data. Our first data release, described here, is the most extensive nuclear phylogenomic dataset for angiosperms to date, comprising 3,099 samples validated by DNA barcode and phylogenetic tests, representing all 64 orders, 404 families (96%) and 2,333 genera (17%). Using the multi-species coalescent, we inferred a “first pass” angiosperm tree of life from the data, which totalled 824,878 sequences, 489,086,049 base pairs, and 532,260 alignment columns. The tree is strongly supported and highly congruent with existing taxonomy, while challenging numerous hypothesized relationships among orders and placing many genera for the first time. The validated dataset, species tree and all intermediates are openly accessible via the Kew Tree of Life Explorer. This major milestone towards a complete tree of life for all flowering plant species opens doors to a highly integrated future for angiosperm phylogenomics through the systematic sequencing of standardised nuclear markers. Our approach has the potential to serve as a much-needed bridge between the growing movement to sequence the genomes of all life on Earth and the vast phylogenomic potential of the world’s natural history collections.

## INTRODUCTION

Discovering the tree of life is among the most fundamental of the grand challenges in science today (Hinchliff et al. 2015). The tree of life is the biological roadmap that allows us to discover, identify and classify life on Earth, to explore its properties, to understand its origins and evolution, and to predict how it will respond to future environmental change. Of all eukaryotic lineages, the angiosperms (flowering plants) are among the most pressing priorities for tree of life research. Angiosperms sustain the terrestrial living world, including humanity, as primary producers, ecosystem engineers and earth system regulators. They hold potential solutions to global challenges, such as climate change, biodiversity loss, human health, food security and renewable energy (Antonelli et al. 2020). In light of this, a phylogenetic framework with which to navigate and interpret the species, trait and functional diversity of angiosperms has never been more necessary. However, despite substantial progress, the evolutionary connections among Earth’s ca. 330,000 flowering plant species (WCVP 2020) remain incompletely known.

The angiosperm research community were early and organised adopters of the molecular phylogenetic approach, resulting in numerous benchmark tree of life publications (e.g. Chase et al. 1993; Soltis et al. 2008; Soltis et al. 2011), and a community approach to phylogenetic classification (APG 1998; APG II 2003; APG III 2009; APG IV 2016). Through this distributed effort, a wealth of DNA sequence data is now available in public repositories, covering ca. 107,000 (31%) of the ca. 350,000 species of vascular plants (RBG Kew 2016; WCVP 2020), most of which are angiosperms (see also Cornwell et al. 2019). However, the lack of sequence data for the remaining 69% obstructs their accurate placement in the tree of life. In addition, lack of complementarity in gene sampling across public DNA sequence data impedes phylogenetic synthesis (Hinchliff and Smith 2014). For example, data from either one or both of *rbcL* and *matK*, the two most popular plastid genes for phylogenetics, are available for only 54% of the ca. 107,000 sequenced vascular plant species (RBG Kew 2016). Comprehensive phylogenetic trees of flowering plants are in high demand (Hinchliff et al. 2015; Eiserhardt et al. 2018), but currently can only be made “complete” using proxies, such as taxonomic classification, to interpolate the unsequenced species (Smith and Brown 2018), which may not accurately reflect relationships. Greater community-wide coordination of both taxon and gene sampling would benefit phylogenetic data integration immensely, creating numerous downstream scientific opportunities.

High-throughput sequencing (HTS) now promises to significantly deepen our understanding of evolutionary relationships among Earth’s species, including angiosperms (Li et al. 2019; Yang et al. 2020). For example, the One Thousand Plant Transcriptomes (1KP) initiative has brought an unprecedented scale of data to bear on the plant tree of life (Wickett et al. 2014; Gitzendanner et al. 2018; Leebens-Mack et al. 2019). Nevertheless, with greatly increased data depth come trade-offs in taxon sampling; the pre-eminent HTS studies cited here account for less than 0.01% of angiosperm species. Undeterred by this sampling gap, the Earth Biogenome Project (EBP) has launched a “moonshot for biology” by proposing to sequence and characterise the genomes of all of Earth’s eukaryotic species over a 10 year period (Lewin et al. 2018). Projects such as the 10,000 Plant Genomes Project (Cheng et al. 2018) and the Darwin Tree of Life Project (https://www.darwintreeoflife.org/) aim to contribute to this goal by producing numerous chromosome-level genome assemblies across major lineages and regional biotas. However, taxon sampling remains a significant issue, due to the challenges of obtaining the high molecular weight DNA required by these projects (for long-read HTS) from samples that are both authentically identified and compliant with the spirit and letter of the Nagoya Protocol (Secretariat of the Convention on Biological Diversity 2011). Despite its immense potential, the “whole genome” approach to discovering the tree of life remains a future goal that will not be achieved on a large taxonomic scale in the short term. Methodological compromises are required to accelerate progress.

The world’s natural history collections are a goldmine for genomic research (Buerki and Baker 2016), containing tissues of almost all species of life on Earth known to science. However, the condition of these tissues and the DNA therein varies widely, depending on age and preservation techniques, among other factors. In the case of plants, herbarium specimens generally yield degraded DNA, which, though not useful for long-read HTS, is now being intensively exploited for short-read HTS (Bakker et al. 2016; Brewer et al. 2019; Forrest et al. 2019; Alsos et al. 2020). In this context, target sequence capture is growing in popularity as the HTS method most widely applied to herbarium DNA (Dodsworth et al. 2019). This approach (also known as target enrichment, target capture, sequence capture, anchored hybrid enrichment) and its variations (e.g. Hyb-Seq, which combines target sequence capture with genome skimming) use RNA or DNA probes to enrich sequencing libraries for specifically targeted loci (Faircloth et al. 2012; Lemmon et al. 2012; Weitemier et al. 2014). It is proving to be an increasingly cost-effective means of isolating hundreds of loci for phylogenetic analysis from even centuries-old specimens (Brewer et al. 2019), bringing comprehensive taxon sampling from herbarium collections within the reach of any phylogenomic researcher (Hale et al. 2020).

Numerous target sequence probe sets have been developed for specific angiosperm groups (e.g. Annonaceae [Couvreur et al. 2019], Asteraceae [Mandel et al. 2014], *Dioscorea* [Soto Gomez et al. 2019], *Euphorbia* [Villaverde et al. 2018]). The design of these probe sets is informed by available genomic resources, as well as criteria specific to the group of interest and research questions. As a result, locus overlap between probe sets tends to be minimal. Unlike the Sanger sequencing era, in which researchers converged on tractable genes such as *rbcL* and *matK*, the lack of complementarity between probe sets curtails prospects for data integration across broad taxonomic scales. In addition, development of custom probe sets is expensive, requiring considerable genomic resources and bioinformatic expertise. A publicly available, universal probe set for angiosperms targeting a standard set of loci would resolve these issues (Buddenhagen et al. 2016; Chau et al. 2018). In response to this, we designed the Angiosperms353 probe set (Johnson et al. 2019), drawing on 1KP transcriptome data from ca. 650 angiosperm species (Leebens-Mack et al. 2019). The probe set targets 353 genes from 410 low-copy, protein-coding nuclear orthologs previously selected for phylogenetic analysis across green plants (Leebens-Mack et al. 2019), enriching up to ca. 260 kbp from any flowering plant. Angiosperms353 probes are an open data resource that can be used without the expense of design or access to prior genomic data and have already been successfully applied across different taxonomic scales (e.g. Larridon et al. 2019; Murphy et al. 2020; Pérez-Escobar et al. 2020; Shee et al. 2020), including at the population level (Van Andel et al. 2019; Slimp et al. 2020; Beck et al. 2021).

Here, we describe a large-scale effort to establish a new phylogenomic platform for exploring the angiosperm tree of life, comprising a set of open tools (Angiosperms353 probes, laboratory protocols, analysis pipeline, data portal) and data (sequence data, assembled genes, alignments, gene trees, species tree). This platform, which directly addresses the challenges outlined above, is an outcome of the Plant and Fungal Trees of Life project (PAFTOL; www.paftol.org) at the Royal Botanic Gardens, Kew (RBG Kew 2015). As a step towards the ultimate goal of a complete species-level tree, we aim to gather DNA sequence data for the Angiosperms353 genes from one species of all 13,862 angiosperm genera (WCVP 2020). This unprecedented dataset of standard loci draws extensively on herbarium collections for comprehensive sampling, especially of genera that have not been sequenced before (Brewer et al. 2019). Extensive new data have been generated, analysed and released into the public domain, along with corresponding phylogenetic inferences. By providing our data in open and accessible ways, including an interactive tree of life, we aim to foster a transparent and collaborative environment for future data re-use and synthesis. This paper serves as the baseline reference for our platform, (i) documenting our methods, (ii) describing our first data release, comprising 17% of angiosperm genera, including initial insights on phylogenetic performance, and (iii) presenting a novel data portal, the Kew Tree of Life Explorer, through which our data and corresponding tree of life can be interrogated and downloaded. We conclude with reflections on the prospects for our approach, future development requirements and the role of open data for enhancing cross-community collaboration towards a complete tree of life.

## MATERIALS AND METHODS

This section describes the workflow (Fig. 1) used by the PAFTOL project to generate our first full data release (i.e. Data Release 1.0), which is publicly accessible through our open data portal, the Kew Tree of Life Explorer (https://treeoflife.kew.org), described below. The workflow consists of three main stages: (i) sample processing, encompassing sample selection and laboratory protocols for target sequence capture data generation (Fig. 2), (ii) data analysis, including target gene assembly, data mining, data validation and phylogenetic inference (Figs. 3, 4), and (iii) data publication via the data portal (Fig. 5). The data accessible via the portal comprise raw data (unprocessed sequence reads) and results from “first pass” analyses (gene assemblies, alignments, gene trees, species tree). Though not exhaustive, these first explorations of the data apply methods that are both rigorous and tractable at our scale of operation.

**Figure 1.**
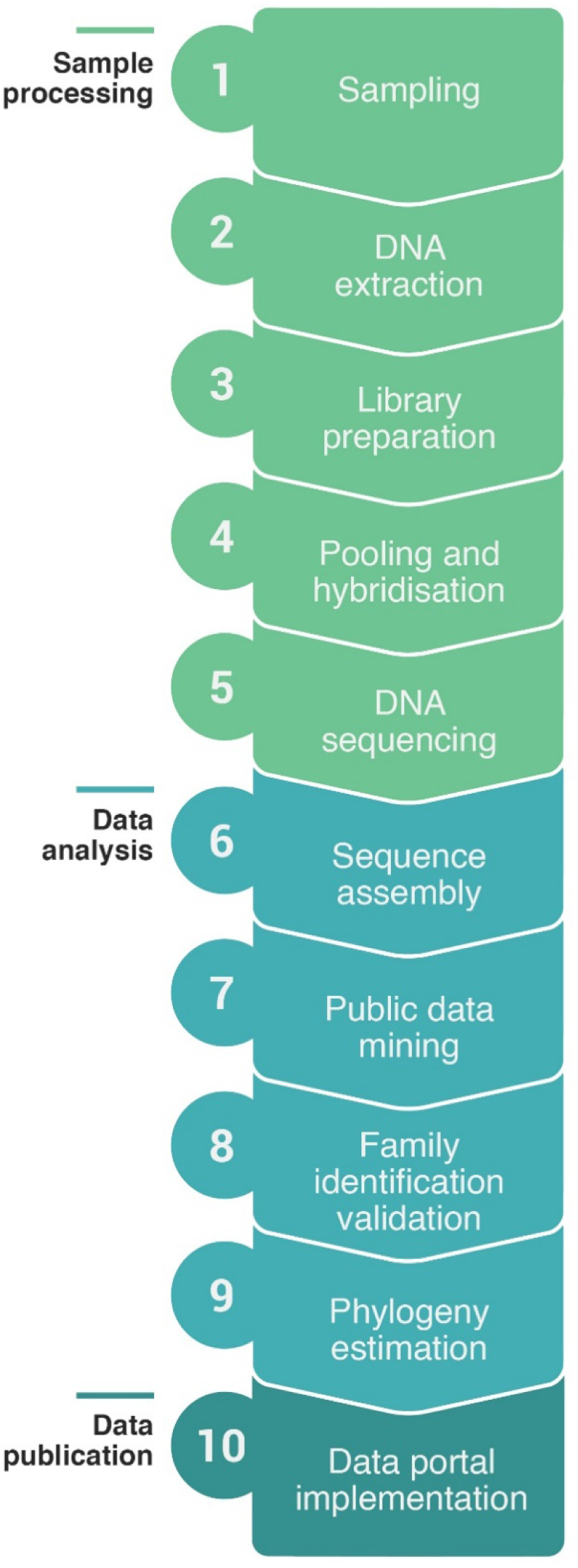
Summary workflow. Overview of steps taken by the PAFTOL project to generate Data Release 1.0 of the Kew Tree of Life Explorer (https://treeoflife.kew.org).

**Figure 2.**
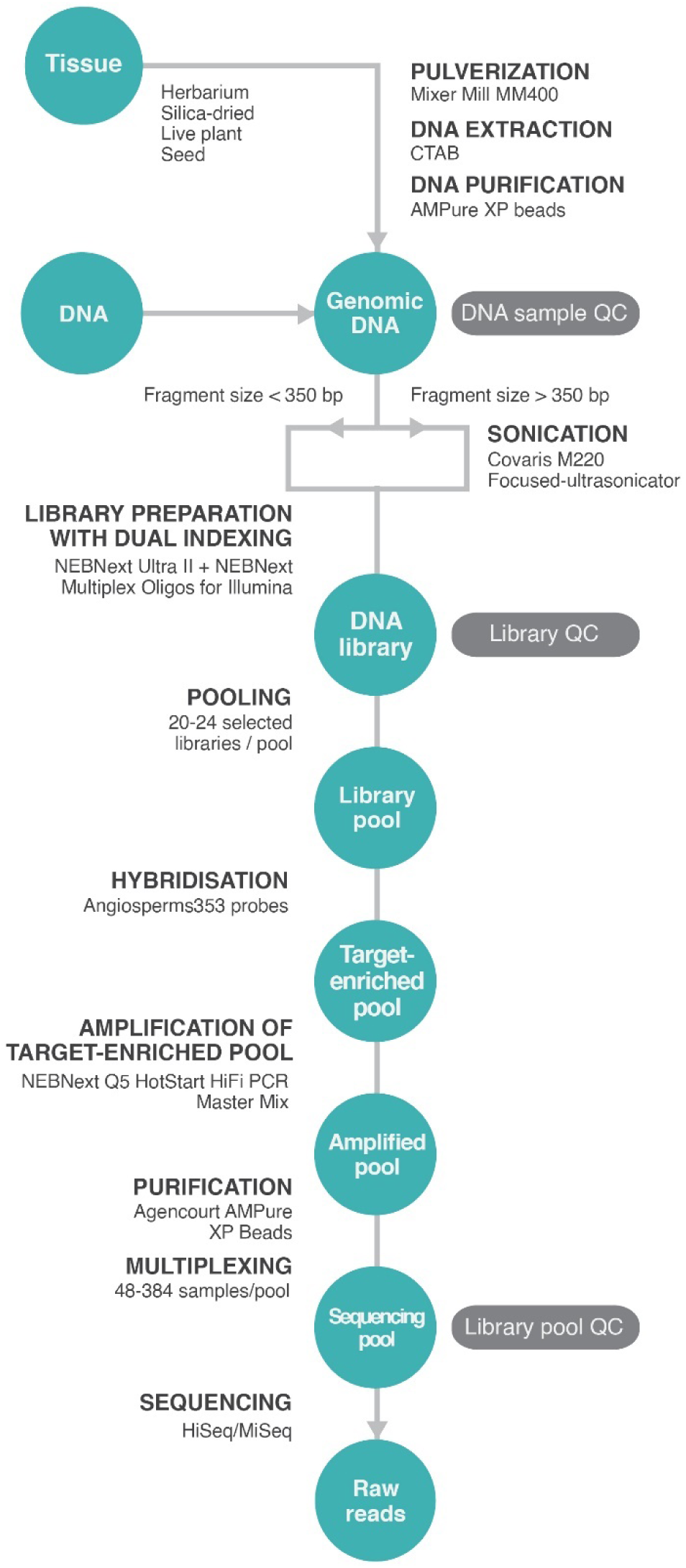
Sample processing workflow. Processes are indicated by bold headings with reagents and machines used given below. Quality control checkpoints are indicated in dark grey boxes.

Details of the first data release are also given in the data release notes in the portal via our secure FTP (http://sftp.kew.org/pub/treeoflife/) and are also archived at the Royal Botanic Gardens, Kew (RBGK) Research Repository (https://doi.org/10.34885/paftol). A new release note will be published in the same locations with each future data release and will detail any changes in methods used relative to the first release described here.

### Sampling

We aimed to generate novel data from across the angiosperms, using a stratified sampling approach of one species per genus. Our sampling was standardised to the complete list of angiosperms within the World Checklist of Vascular Plants (WCVP 2020), which currently recognises 13,862 accepted genera in 418 families, aligned to the 64 orders of the APG IV classification (APG IV 2016). We prioritised genera that were not represented by published transcriptomic or genomic data in public sequence repositories (e.g. GenBank), and avoided genera that had already been sampled in large genomic initiatives such as the 1KP project (Leebens-Mack et al. 2019). The selection of species within genera was made pragmatically, although we prioritised the species of the generic type where possible.

Plant material was obtained from a variety of sources (Fig. 2), primarily from the collections of RBGK (herbarium, DNA bank, silica gel-dried tissue collection, living collection and the Millennium Seed Bank, https://www.kew.org/science/collections-and-resources/collections). Additional material (tissue samples, extracted DNA) was generously provided by our collaborative networks (see Acknowledgements). To be selected, the material must have been (i) legally sourced and made available for use in phylogenomic studies, (ii) identified to species level, preferably by an expert of the group, and (iii) ideally collected in the wild. As far as was practically achievable, we ensured that the identity of each sample was substantiated by a voucher specimen deposited in a publicly accessible herbarium.

All metadata were captured using a relational database that allowed us to track processing of samples from the selection of material, through the library preparation pipeline to the completion of sequencing. Data were recorded in four main tables (Specimen, Sample, Library, Sequencing). The database architecture allowed us to record multiple sequence datasets (fastq files) from one or several libraries, and one or several DNA extracts from a single specimen. Relevant voucher specimen information was also captured in the database (e.g. collector(s), collector number, herbarium acronym (following Index Herbariorum http://sweetgum.nybg.org/science/ih/), country of origin, date of collection, specimen barcodes). Voucher data are available via our data portal (see below). Images of specimens sampled from the RBGK Herbarium are in the process of being captured in RBGK’s online herbarium catalogue (http://apps.kew.org/herbcat/) and, where available, are linked to the appropriate records in the Kew Tree of Life Explorer.

### DNA extraction

DNA was extracted from 40 mg of herbarium material, 20 mg of silica gel-dried material (Chase and Hills 1991), or 100 mg of fresh material using a modified CTAB extraction method (Doyle and Doyle 1987; Fig. 2). Plant tissue was pulverized using a Mixer Mill MM400 (Retsch GmbH, Germany). DNA extractions were purified by a magnetic bead clean-up using Agencourt AMPure XP beads (Beckman Coulter, Indianapolis, IN, USA), according to the manufacturer’s protocols. Samples obtained from the RBGK DNA bank (http://dnabank.science.kew.org/homepage.html) had been extracted using a modified CTAB method (Doyle and Doyle 1987) followed by caesium chloride/ethidium bromide density gradient cleaning and dialysis. DNA samples provided by external collaborators had been extracted using a wide variety of extraction methods from living, silica gel-dried and herbarium material. All DNA samples were quality checked for concentration and degree of fragmentation. DNA concentration was measured using a Quantus (Promega, Madison, WI, USA) or Qubit (Thermo Fisher Scientific, Inchinnan, UK) fluorometer. DNA fragment size range was routinely assessed on a 1% agarose gel using ethidium bromide and visualized with a UVP Gel Studio (AnalytikJena, Jena, Germany). For samples with a low DNA concentration (i.e. not visible on a gel), fragment sizes were assessed on a 4200 TapeStation using Genomic DNA ScreenTape (Agilent Technologies, Cheadle, UK).

### Library preparation

Genomic DNA samples were diluted to 4 ng/µl with 10 mM Tris (pH 8.0). Those with an average fragment size greater than 350 bp were sonicated to an average fragment size ca. 400 bp, using a Covaris M220 Focused-ultrasonicator (Covaris, Woburn, MA, USA) by adding 50 µl of diluted genomic DNA to a 130 µl Covaris microAFA tube. The sonication time was adjusted for each sample based on its average DNA fragment size (15 to 100 secs, following the manufacturer’s protocols). Additional parameters used were peak incident power to 50W, duty factor to 10% and 200 cycles per burst.

Libraries were prepared using the NEBNext Ultra II DNA Library Prep Kit (New England Biolabs, Ipswich, MA, USA; Fig. 2). Size selection was not employed for samples with highly degraded DNA. In the early stages of the project, libraries were prepared following the manufacturer’s protocols exactly, but the majority were prepared using half of the recommended volumes throughout to increase cost efficiency. All DNA fragments were indexed using NEBNext Multiplex Oligos for Illumina (Dual Index Primer sets 1 and 2, New England Biolabs, Ipswich, MA, USA).

The distribution of fragment sizes in each library was assessed with a 4200 TapeStation using standard D1000 tapes. Library concentration was measured using a Quantus fluorometer. If the library concentration was less than 10 nM, up to eight additional PCR cycles were performed, following the NEBNext Ultra II Library Prep Kit protocol with IS5_reamp.P5 and IS6_reamp.P7 primers (Meyer and Kircher 2010). Library quality assessment was then repeated.

### Pooling and hybridisation

Prior to hybridisation (Fig. 2), all libraries were normalised to 10 nM, using 10 mM Tris (pH 8.0) and then combined into pools of 20 to 24 libraries, each containing 10 µl (0.1 pmol) of each normalized library (i.e. a total of ca. 600-700 ng DNA in each pool, assuming an average fragment size of ca. 450 bp). To ensure even sequencing across all samples in a pool, species for pooling were selected to minimize the range of DNA fragment sizes and ensure a narrow taxonomic breadth. The latter criterion was needed because samples that are more closely related to the taxa used to construct the probe set tend to preferentially hybridise. This can lead to an over-representation of their sequences in the DNA data if appropriate care is not taken when selecting species for the sequencing pool. In rare cases, such as smaller pools (ca. 10 libraries) of short fragment (i.e. <300 bp) libraries, it was necessary to recalculate the standard volume of normalized libraries to be added to ensure that the final pool contained ca. 500 ng of DNA.

The pooled libraries were dried in a SpinVac (Eppendorf, Dusseldorf, Germany), resuspended in 8 µl of 10 mM Tris (pH 8.0) and enriched by hybridising with the Angiosperms353 probe kit (Johnson et al. 2019; Arbor Biosciences myBaits Target Sequence Capture Kit, ‘Angiosperms 353 v1’, Catalogue #308196) following the manufacturer’s protocol, version 4.0. Hybridisation was typically performed at 65°C for 24 h, with reactions topped with 30 μl of red Chill-out Liquid Wax (Bio-Rad, Hercules, CA, USA) to prevent evaporation. However, for short libraries (i.e. <350 bp) the temperature was reduced to 60°C, following the recommendations of Arbor Biosciences.

The target-enriched pools were amplified using the KAPA HiFi 2X HotStart ReadyMix PCR Kit (Roche, Basel, Switzerland) or NEBNext Q5 HotStart HiFi PCR Master Mix (New England BioLabs, Ipswich, MA, USA) for eight to 14 cycles. Amplified pools were then purified using Agencourt AMPure XP Beads (at 0.9X the sample volume) and eluted in 15 µl of 10 mM Tris (pH 8.0).

Products were quantified with a Quantus fluorometer and re-amplified if the concentration was below 6 nM, with three to six PCR cycles (see above). Final products were assessed using the TapeStation to determine the distribution of fragment sizes. The target-enriched pools were normalized to 6 nM (using 10 nM Tris, pH 8.0) and multiplexed for sequencing, with the number of target-enriched pools combined in each sequencing pool varying from two to 20 (comprising a total of 48-384 samples) depending on the sequencing platform and service provider requirements.

### DNA sequencing

Initially, DNA sequencing was performed on an Illumina MiSeq at RBGK with version 3 chemistry (Illumina, San Diego, CA, USA) and ran for 600 cycles to generate 2 × 300 bp paired-end reads. Subsequently, DNA sequencing was outsourced (Macrogen, Seoul, South Korea, or Genewiz, Takeley, UK) and performed on an Illumina HiSeq producing 2 × 150 bp paired-end reads. Raw reads were deposited in the European Nucleotide Archive under an umbrella project (accession number PRJEB35285) and can be accessed from the individual sample records in the Kew Tree of Life Explorer.

### Sequence assembly

Coding sequences were recovered from target-enriched sequence data using our pipeline recoverSeqs (accessible from our GitHub repository https://github.com/RBGKew/KewTreeOfLife, pypaftol ‘paftools’ submodule) to retrieve sequences orthologous to the Angiosperms353 target gene set (Johnson et al. 2019; https://github.com/mossmatters/Angiosperms353). This target set contained multiple reference sequences per gene, thereby covering a large phylogenetic breadth to facilitate read recovery across angiosperms.

The process comprised four main stages (Fig. 3), applied to each sample: (i) sequence reads were trimmed using Trimmomatic (Bolger et al. 2014) with the following parameters: ILLUMINACLIP: <ADAPTERFASTAFILE>: 2:30:10:2:true, LEADING: 10, TRAILING: 10, SLIDINGWINDOW: 4:20, MINLEN: 40, with the adaptor fasta file formatted for palindrome trimming, (ii) trimmed read pairs were mapped to the Angiosperms353 target genes with TBLASTN. A representative reference sequence for each gene was then selected by identifying the sequence with the largest number of mapped reads. (iii) This representative gene was used as the reference for assembling the gene-specific reads using an overlap-based assembly algorithm (--assembler overlapSerial option) as follows. First, the reads were aligned to and ordered along the reference sequence based on a minimum alignment size of 50 bases (--windowSizeReference option) with a minimum sequence identity of 70% (--relIdentityThresholdReference option). Consecutive reads ordered along the reference sequence were aligned in a pair-wise manner to find read overlaps. If an overlap of at least 30 bases (--windowSizeReadOverlap option) and 90% sequence identity (--relIdentityThresholdReadOverlap option) was found, the aligned reads were used to construct a consensus contig with ambiguous bases represented by ‘N’. This last parameter resulted in one or more sets of aligned reads with ≥90% sequence identity, each set being merged into a single contig. In the final stage, the exonerate protein2genome program was used to identify the exon-intron structure within each contig. One or more contigs were chosen that best represented the structure of the exon(s) in the reference gene chosen in step (ii). If the exons existed in multiple contigs, those contigs were joined together to form the recovered gene coding sequence.

**Figure 3.**
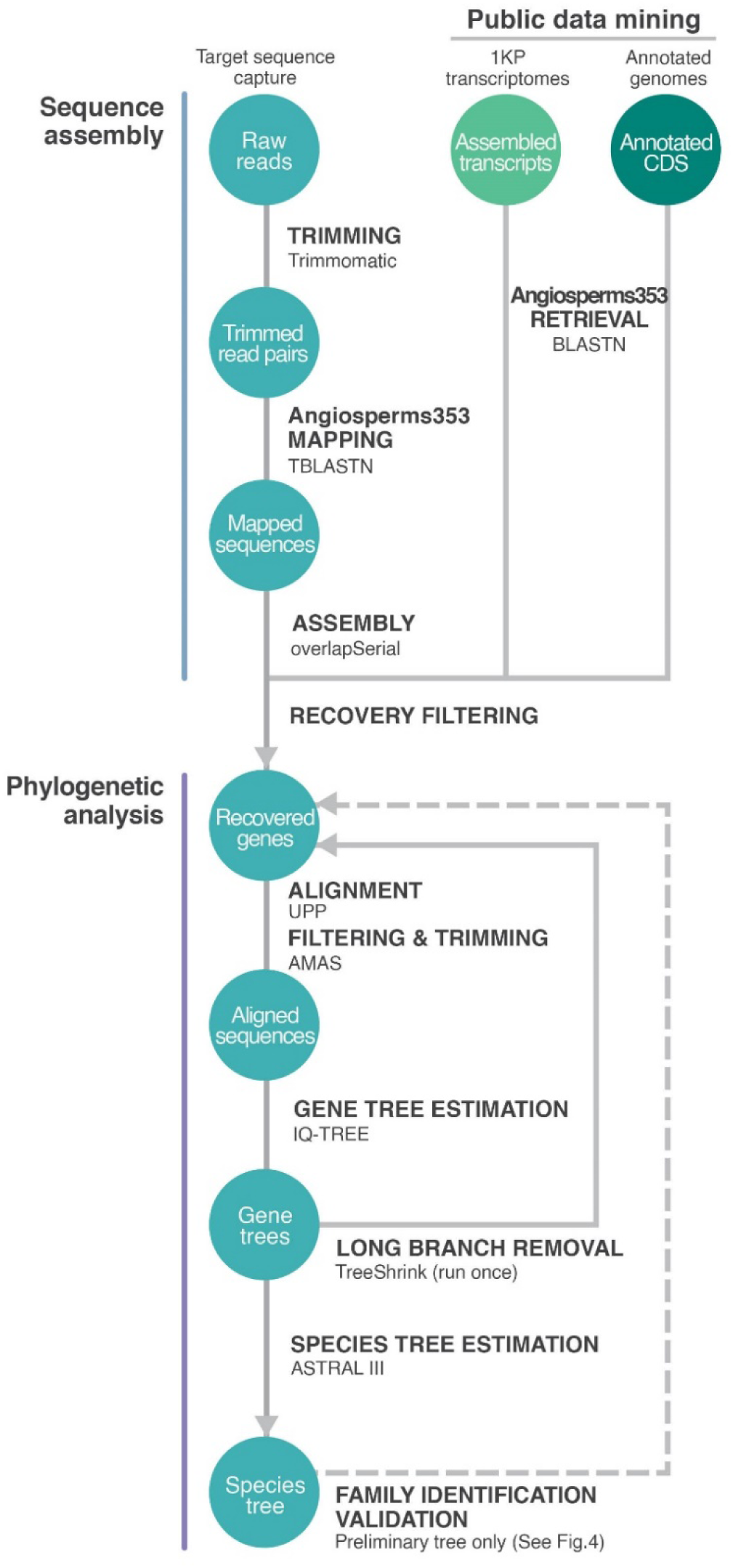
Data analysis workflow. Pipeline products are shown in blue-green circles (available to download via the Kew Tree of Life Explorer, https://treeoflife.kew.org). Processes are indicated by bold headings with programs used given below.

Target gene recovery success was assessed for each sample by calculating the number of genes recovered and the sum of the recovered gene lengths. Samples were removed from downstream analyses if the sum of the recovered gene lengths fell below 20% of the median value across all samples.

### Public data mining

In addition to newly generated target sequence capture data, the Angiosperms353 genes were mined from publicly available genomic data (Fig. 3). For the first release, we mined data from the 1KP Initiative (Carpenter et al. 2019; Leebens-Mack et al. 2019) and published genomes with gene annotations (https://plants.ensembl.org/). The genes were retrieved from assembled transcript sequences (1KP) or coding sequences (CDS; genomes) using paftools retrievetargets from our pipeline, which relies on BLASTN to identify and extract the genomic or transcriptomic sequences corresponding to the 353 genes. Because initial recovery of genes from 1KP transcripts was unsatisfactory, we expanded the Angiosperms353 target set (dataset available from our GitHub) to improve matching and retrieval of genes. As with the novel target sequence capture assemblies, data were removed from downstream analyses if the sum of the gene lengths fell below 20% of the median value across all samples.

### Family identification validation

To verify the family identification of our processed samples, we implemented two validation steps, which were run in parallel (Fig. 4). The two steps consisted of (i) DNA barcode validation, which utilised nuclear ribosomal and plastid barcodes for DNA-based identification, and (ii) phylogenetic validation, which checked the placement of each sample in a preliminary tree relative to its expected position based on its initial family assignment. Identification checks below the family level were not conducted due to the incompleteness of adequate reference resources for DNA barcode validation and sparseness of sampling for phylogenetic validation at the genus or species level.

**Figure 4.**
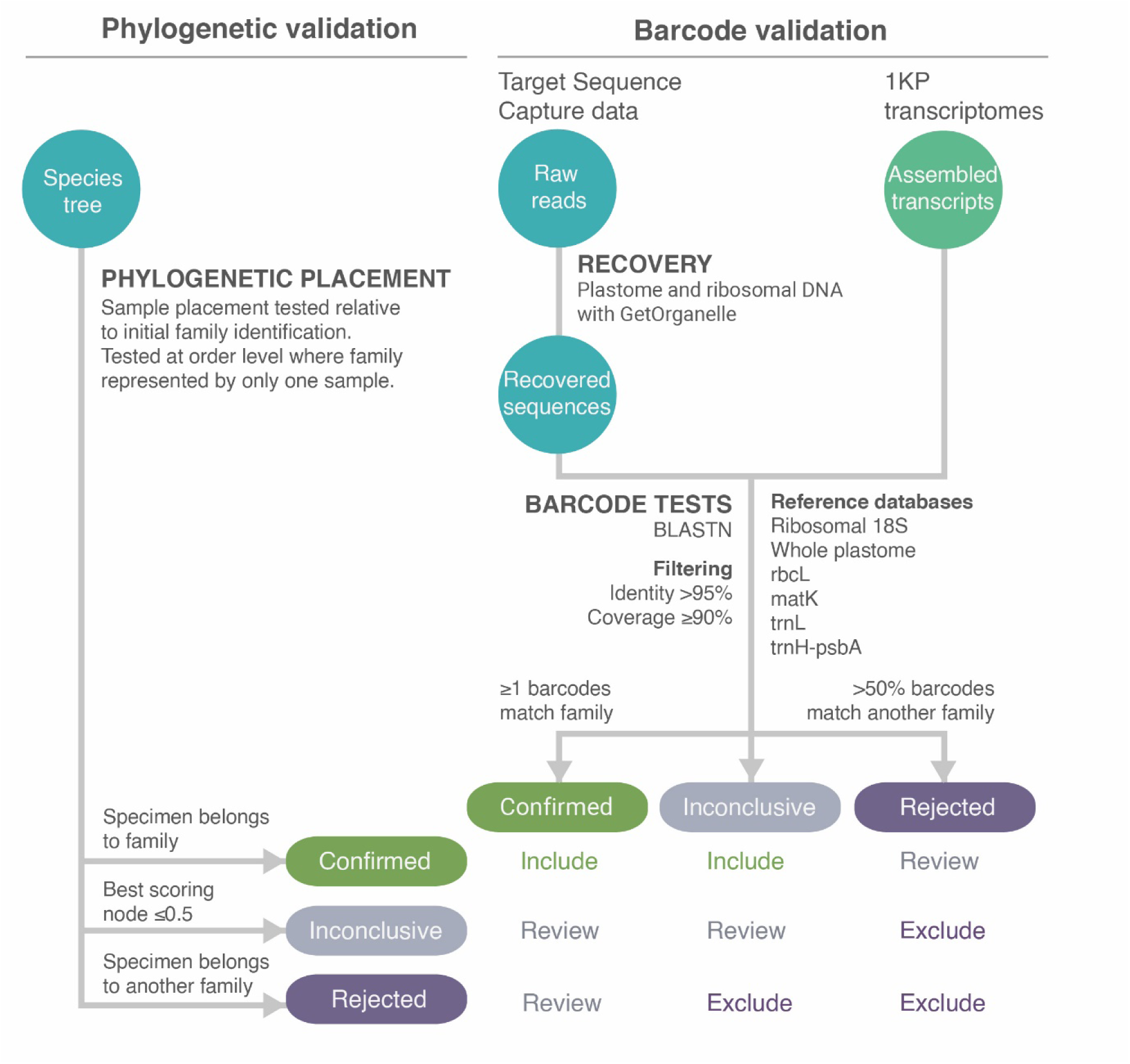
Family identification validation workflow. Processes are indicated by bold headings. Embedded table (bottom right) indicates decisions made for each sample based on the two validation steps.

For barcode validation of target sequence capture data (Fig. 4), plastomes and ribosomal DNA were recovered from raw reads using GetOrganelle (Jin et al. 2020) and subsequently queried against databases of reference plant barcodes using BLASTN (Camacho et al. 2009). For 1KP samples, transcriptome assemblies were directly used as queries in BLASTN. Note that we considered the family identity of annotated genomes to be correct and hence a barcode validation was unnecessary. Six individual barcode reference databases were built from the NCBI nucleotide and BOLD databases (https://www.ncbi.nlm.nih.gov/nuccore; https://www.boldsystems.org/, accessed on 29/10/2020), one for the whole plastome, and the remaining five for specific loci (nuclear ribosomal 18S, as well as plastid *rbcL*, *matK*, *trnL*, and *trnH-psbA*). As for samples, the taxonomy of reference sequences was standardized to WCVP (WCVP 2020). BLAST results were further filtered with a minimum identity >95% and a minimum coverage of reference locus ≥90% (except for whole plastomes, for which only a filtering based on minimum length was applied).

Tests could only be completed if a sample’s given family was present in the barcode databases and if at least one BLAST match remained after filtering. Thus, zero to six barcode tests were conducted per sample. A sample passed an individual test if the first ranked BLAST match (ranked by percentage of identity) confirmed its original family identification and failed otherwise. The final result of the barcode validation following the six individual barcode tests were determined as follows: (i) Confirmed, if one or more barcode tests matched the family identification of a sample; (ii) Rejected, if more than half of the barcode tests gave the same incorrect family identification (requires at least two barcode tests); (iii) Inconclusive (otherwise). Further details of the barcode validation methods can be found in Supplementary Material available on Dryad. The scripts and lists of NCBI and BOLD accessions used in barcode databases are available on our GitHub repository.

To conduct phylogenetic validation (Fig. 4), a preliminary phylogenetic tree was built using the complete, unvalidated dataset, following the phylogenetic methods described below. We then assessed which nodes best represented each order and family in the tree. For every node in the tree, two metrics were calculated for all families and orders: (i) the proportion of samples belonging to a given order/family that are descendants of the node, and (ii) the proportion of samples descending from the node that belong to the order/family. The two metrics were then multiplied to produce an overall taxon concordance score. For each family and order, the highest scoring node was subsequently considered to best represent the taxon in the tree (allowing the identification of outlying samples). A node with a score of 1 for a given order/family is the crown node (most recent common ancestral node) of that taxon, which is monophyletic in the tree. See Supplementary Figure S1 for an illustration.

The family identification of each sample was determined as (i) Confirmed: if identified as belonging to a family whose best scoring node had a taxon concordance score >0.5 and found as a descendant of this node in the tree, (ii) Rejected: if identified as belonging to a family whose best scoring node had a taxon concordance score >0.5 but not found as a descendant of this node, or (iii) Inconclusive: if identified as belonging to a family whose best scoring node had a taxon concordance score ≤0.5. Note that for families represented in the tree by a single sample, the validation was performed with respect to their orders. If the order was represented by a single sample, the validation result was coded as inconclusive.

The outputs of the phylogenetic and DNA barcode validation were combined to identify samples for automatic inclusion and exclusion from the final dataset, and samples for which a decision on inclusion/exclusion was subject to expert review (Fig. 4). Exclusions after expert review were made based on implausible tree placement (e.g. wrong higher clade) or sample misidentification (e.g. match to another family in the barcode validation).

All assembled Angiosperms353 gene data from all samples validated for inclusion form the basis of Data Release 1.0. These were made publicly available via the Kew Tree of Life Explorer.

### Phylogeny estimation

We inferred a phylogenetic tree from all validated data (Data Release 1.0) for presentation in an interactive format in the Kew Tree of Life Explorer. This species tree was estimated from gene trees using the multi-species coalescent summary method implemented in ASTRAL-III (Zhang et al. 2018). In addition to the angiosperm samples, ten samples representing seven gymnosperm families from the 1KP initiative were mined for Angiosperms353 orthologs and included in all analyses as outgroup taxa. Our phylogenomic pipeline, available from our GitHub repository, is summarised below. For each gene, DNA sequences were aligned with UPP 4.3.12 (Nguyen et al. 2015).

At the start of the alignment process a set of 1,000 sequences were selected for an initial backbone tree. Option -M was set to ‘-1’ so that sequences could be selected within 25% of the median full-length sequence. Filtering and trimming of the alignment were performed with AMAS (Borowiec 2016) as follows. Sequences with insufficient coverage (<60%) across well occupied columns of each gene alignment were removed. Well occupied columns were defined as those with more than 70% of positions occupied. Then, alignment columns with <0.3% occupancy were removed to avoid a large number of columns with very rare or unique insertions from being included in the tree reconstruction. Finally, sequences with a total length of less than 80 bases were removed, and genes with <30 overlapping bases (at the 70% threshold mentioned above) were excluded.

Gene trees were estimated with IQ-TREE 2.0.5 (Minh et al. 2020) inferring branch support using the ultrafast bootstrap method (option -B; Hoang et al. 2017) with the maximum number of iterations set to 1,000 (option -nm) and using a single model of evolution (option -m GTR+F+R). The use of a single model without testing many models of evolution was a pragmatic choice, following Abadi et al. (2019). TreeShrink 1.3.4 (Mai and Mirarab 2018) was used to remove abnormally long branches from gene trees using default settings, except option -b, which was set to 20. The alignment and gene tree estimation steps were then repeated on the samples retained by TreeShrink. Before reconstructing the species tree using ASTRAL-III, nodes in the gene trees with bootstrap support values less than 30% were collapsed using nw_ed from Newick Utilities 1.6.0 (Junier and Zdobnov 2010). This value was deduced from interpreting Figure 1 in Hoang et al. (2017), adjusting the standard bootstrap threshold of 10% (recommended for ASTRAL-III), to 30 % for the ultrafast bootstrap.

All gene alignments, gene trees and the ASTRAL-III species tree are available for download from secure FTP and the Kew Tree of Life Explorer. In addition, the species tree is available to browse through an interactive tree viewer implemented within the Kew Tree of Life Explorer (see also Supplementary Fig. S2).

### Data portal implementation

To disseminate results, a data portal (the Kew Tree of Life Explorer; https://treeoflife.kew.org) was designed and implemented (Fig. 5) with a layered architecture that comprised: (i) a MariaDB running on a Galera multi-master cluster as a database management system; (ii) an API written in Python using the Flask framework and the SQLAlchemy library; (iii) a front-end written using the Vue.js framework and Nuxt.js for the tabular data (used to provide access to gene and specimen data) and content pages; (iv) a tree visualisation module developed from the open source application PhyD3 (Kreft et al. 2017) using D3.js (Bostock 2012) for data visualisation; and (v) deployment on a Linux (CentOS 7) server using Nginx as web server and load balancer.

**Figure 5.**
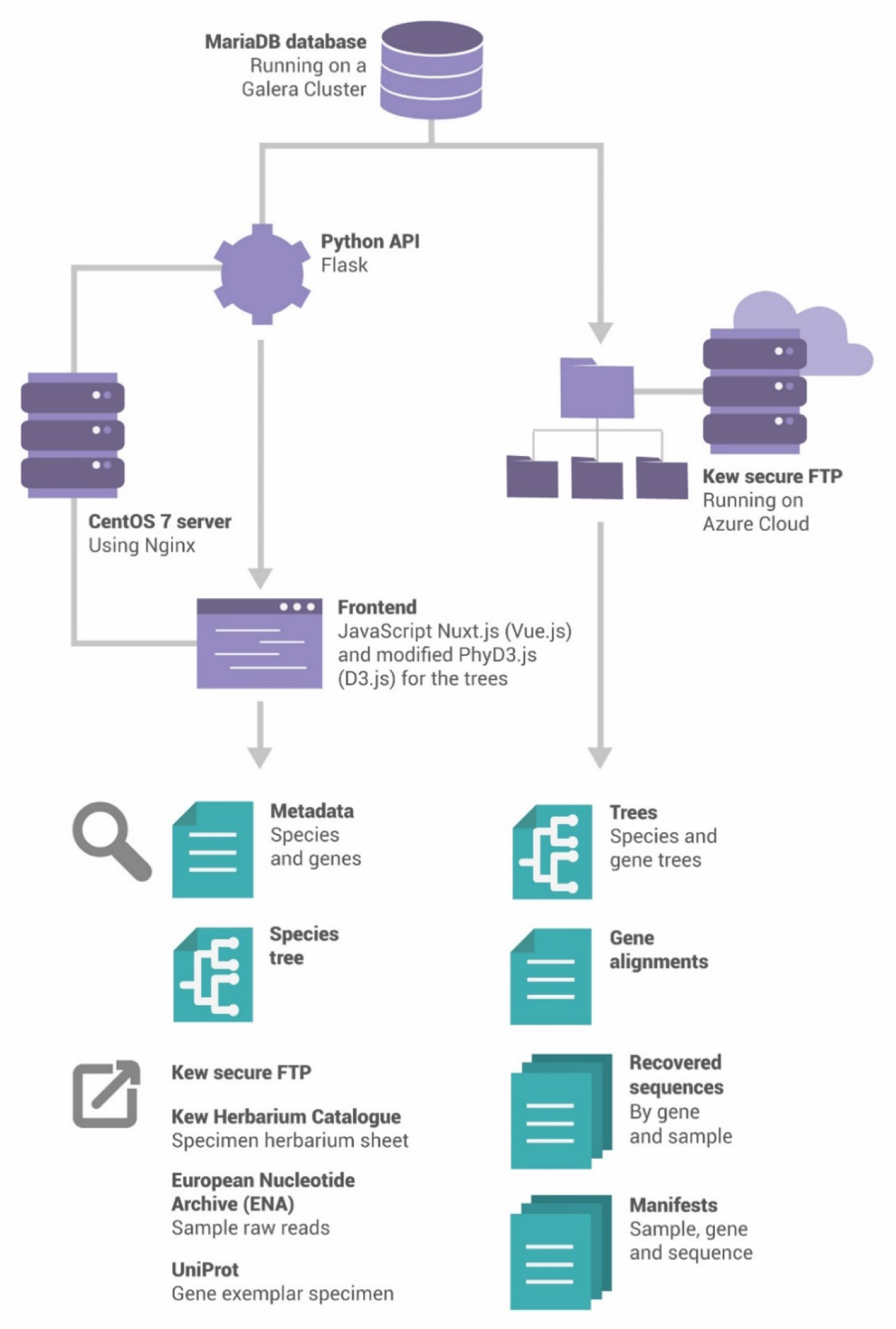
Data publication workflow. Implementation of Kew Tree of Life Explorer data portal is illustrated. Arrows indicate data flow from internal repository to public interface. Infrastructural components are shown in purple; publicly available information is shown in green. External links available from the portal are listed in the lower left.

The data, with appropriate metadata and documentation, are available for public download over secure FTP (http://sftp.kew.org/pub/treeoflife/) and the Kew Tree of Life Explorer under a Creative Commons Attribution 4.0 International (CC BY 4.0) license. When superseded by new releases, archived earlier releases will remain accessible via secure FTP.

## RESULTS

### Initial dataset

The initial dataset prior to processing and analysis comprised data from 3,272 angiosperm samples, representing 413 families of angiosperms (99%) and 2,428 genera (18%; Table 1). We generated novel target sequence capture data for 2,522 of these samples, which included 104 angiosperm genera that have never been sequenced before. Data for the remainder were mined from public sources (689 1KP transcriptomes, 61 annotated genomes). The majority of target sequence capture data were generated from the RBGK collections as follows: DNA Bank (43%), herbarium (28%), silica gel-dried tissue collection (8%), living collection (2%), and Millennium Seed Bank (0.3%). The remaining 19% of samples included in this study were provided by various collaborators of the PAFTOL project, either as DNA samples or as dried tissue (see Acknowledgements).

**Table 1.** Total number of angiosperm samples included at three stages of data release preparation. The first column represents all samples available in the initial dataset. The second column indicates samples included in our preliminary tree, prior to family identification validation, but after removal of samples for which the sum of the gene lengths fell below 20% of the median value across all samples. The third column provides numbers for the samples made public in the Kew Tree of Life Explorer, Data Release 1.0, and included in our final phylogenetic tree. Numbers of angiosperm families, genera and species in each data subset are provided in brackets (as families/genera/species).

| Data source | Initial dataset | Preliminary tree pre-validation | Final tree and Data Release 1.0 |
| --- | --- | --- | --- |
| Target sequence capture data | 2,522<br>(304/1988/2397) | 2,438<br>(297/1947/2340) | 2,374<br>(292/1903/2280) |
| 1KP transcriptomes | 689<br>(254/544/682) | 678<br>(250/530/677) | 664<br>(245/517/663) |
| Annotated genomes | 61<br>(23/43/59) | 61<br>(23/43/59) | 61<br>(23/43/59) |
| **Total** | **3,272**<br>**(413/2428/3079)** | **3,177**<br>**(410/2388/3028)** | **3,099**<br>**(404/2333/2956)** |

Sequence recovery from all 2,522 target sequence capture samples (prior to any quality controls) is visualised in Figure 6. Eighty-four target sequence capture samples and eleven 1KP transcriptomes were removed from downstream analyses because the sum of gene lengths did not meet the quality threshold of 20% of the median value across all samples.

**Figure 6.**
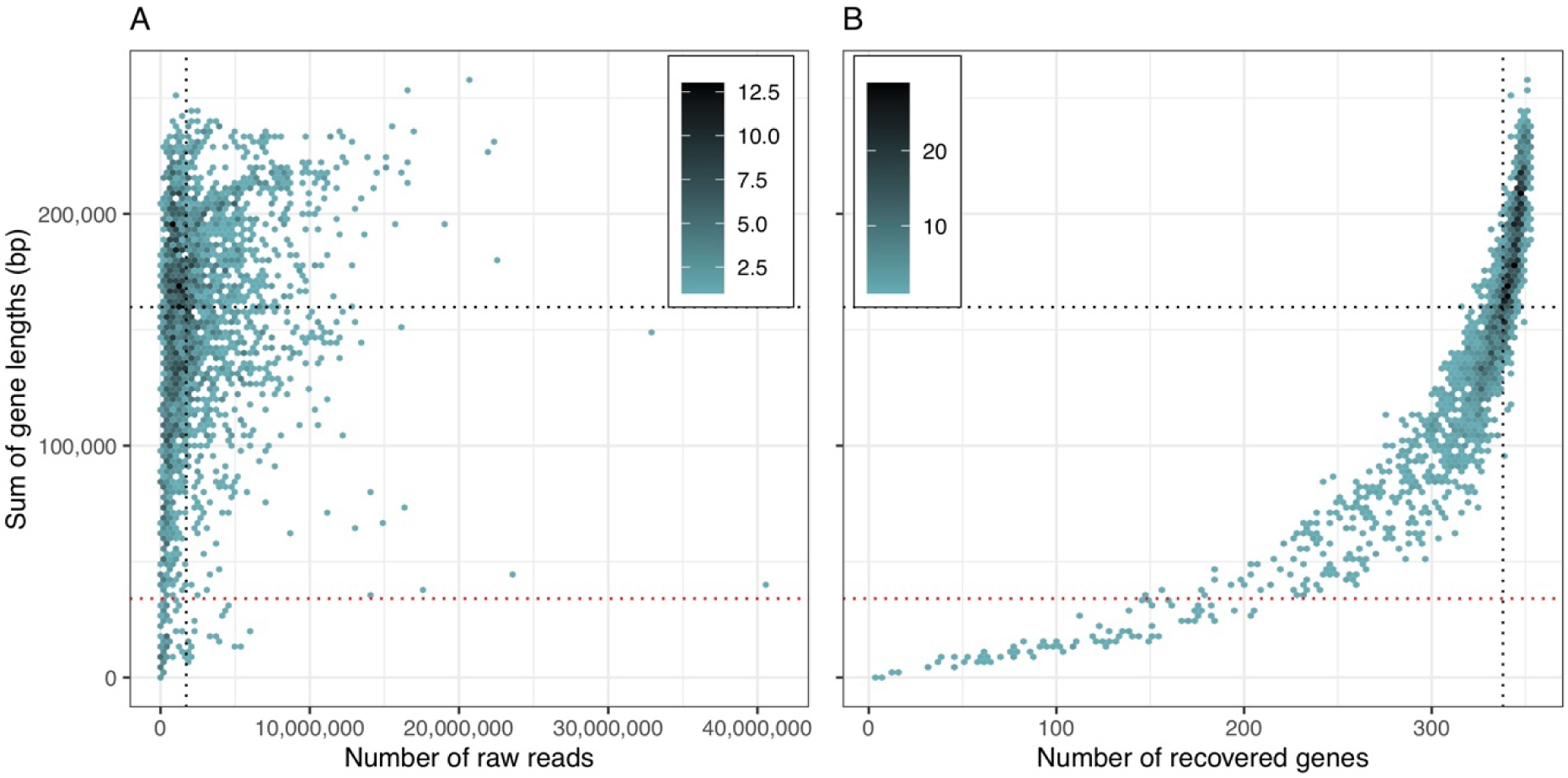
Density plots of target sequence recovery from our raw data. Data are presented prior to any filtering, illustrating relationships of sum of gene lengths (bp) to (a) the number of raw reads and (b) the number of recovered genes. Colours indicate density of data points. Black dotted lines indicate medians of variables and red dotted lines indicate the threshold used to remove samples from downstream analyses, set as 20% of the median value across all samples.

### Family identification validation

The remaining 3,177 samples (Table 1) were processed through our sample family identification validation pipeline (Fig. 4, Table 2, Supplementary Table S1). Of these, 3,064 (97%) were automatically cleared for inclusion and 67 were automatically excluded (Table 2). The remaining 46 samples were held for expert review, after which 35 were cleared for inclusion and 11 were excluded due to implausible tree placements. The majority of excluded samples (64 out of 78) were from the novel target sequence capture data, although 14 were 1KP transcriptomes, highlighting the risk of sample misidentification in even the most highly curated datasets. Further details regarding the results obtained during the family identification validation by DNA barcoding can be found in Supplementary Material available on Dryad.

**Table 2.** Results of validation of sample family identification. The family identification of each sample was scored as confirmed, inconclusive or rejected according to both DNA barcode and phylogenetic validations. Where only a single-family representative was included, samples were tested at the ordinal level. Based on these results, samples were automatically included, excluded, or held for review. See Materials and Methods and Fig. 4 for more details.

|  |  | DNA barcode validation |  |  |
| --- | --- | --- | --- | --- |
|  |  | Confirmed | Inconclusive | Rejected |
| Phylogenetic<br>validation | Confirmed | 2,666 | 398 | 4 |
|  |  | Include | Include | Review |
|  | Inconclusive | 27<sup>a</sup> | 7 | 3 |
|  |  | Review | Review | Exclude |
|  | Rejected | 8 | 42 | 22 |
|  |  | Review | Exclude | Exclude |

<sup>a</sup>Samples with confirmed family (barcode), but for which the placement cannot be confidently assessed were reviewed.

The final validated dataset for Data Release 1.0 consisted of 3,099 angiosperm samples (Table 1), only 5% fewer than were present in the initial dataset. These samples represent all 64 orders, 404 families (96%; 212 represented by >1 sample), 2,333 genera (17%) and 2,956 species (0.01%).

### Data Release 1.0: sequence quality and gene recovery

Nine statistics were used to assess the sequence quality across the 3,099 samples of Data Release 1.0 (Table 3). For the 2,374 target sequence capture samples, the mean percentage of on-target reads was 8%, the mean read depth per sample across all recovered genes was 90x with a median value of 38x and the mean percentage length of recovered genes per sample was 62%. The number of genes and the sum length of gene sequence recovered per sample were tightly correlated as expected, varying continuously across the dataset up to the full set of Angiosperms353 genes and a total gene length of 256.9 kbp, close to the maximum expected length of 260 kbp for recovering genes with this target gene set (Fig. 6). However, both the number of genes and sum length of gene sequence recovered were correlated less closely with the number of available reads than they were to each other. The total length of sequence recovered from target sequence capture data was shorter than for samples mined for Angiosperms353 genes from 1KP transcriptomes or annotated genomes data (Table 3). The reason for the shorter length of the recovered genes is that some exons were absent from the original 1KP alignments used by Johnson et al. (2019) to create the Angiosperms353 gene set. These missing exons are however present in 1KP transcriptomes and annotated genomes and were recovered during data mining. The variation in performance of target enrichment across different samples, illustrated by the measures of variability shown in Table 3, likely reflects the variation in structure and metabolite composition of the starting tissue, which is known to impede DNA extraction from various species and its downstream manipulation. This variation is one of the challenges in dealing with samples from a broad taxonomic range such as across the evolutionary diversity of angiosperms. Variation in gene recovery across orders is visualised in Supplementary Figure S3.

**Table 3.**
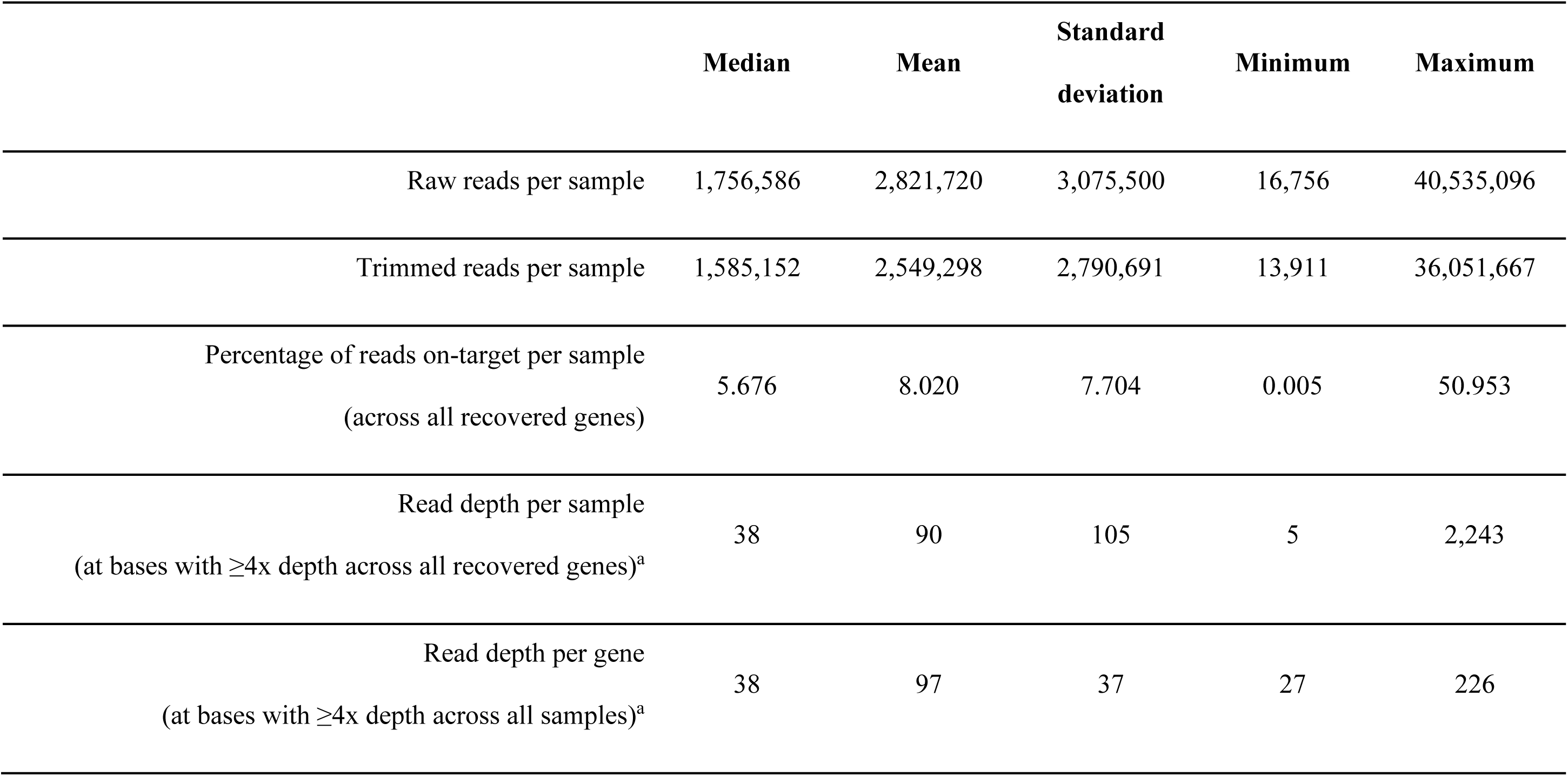
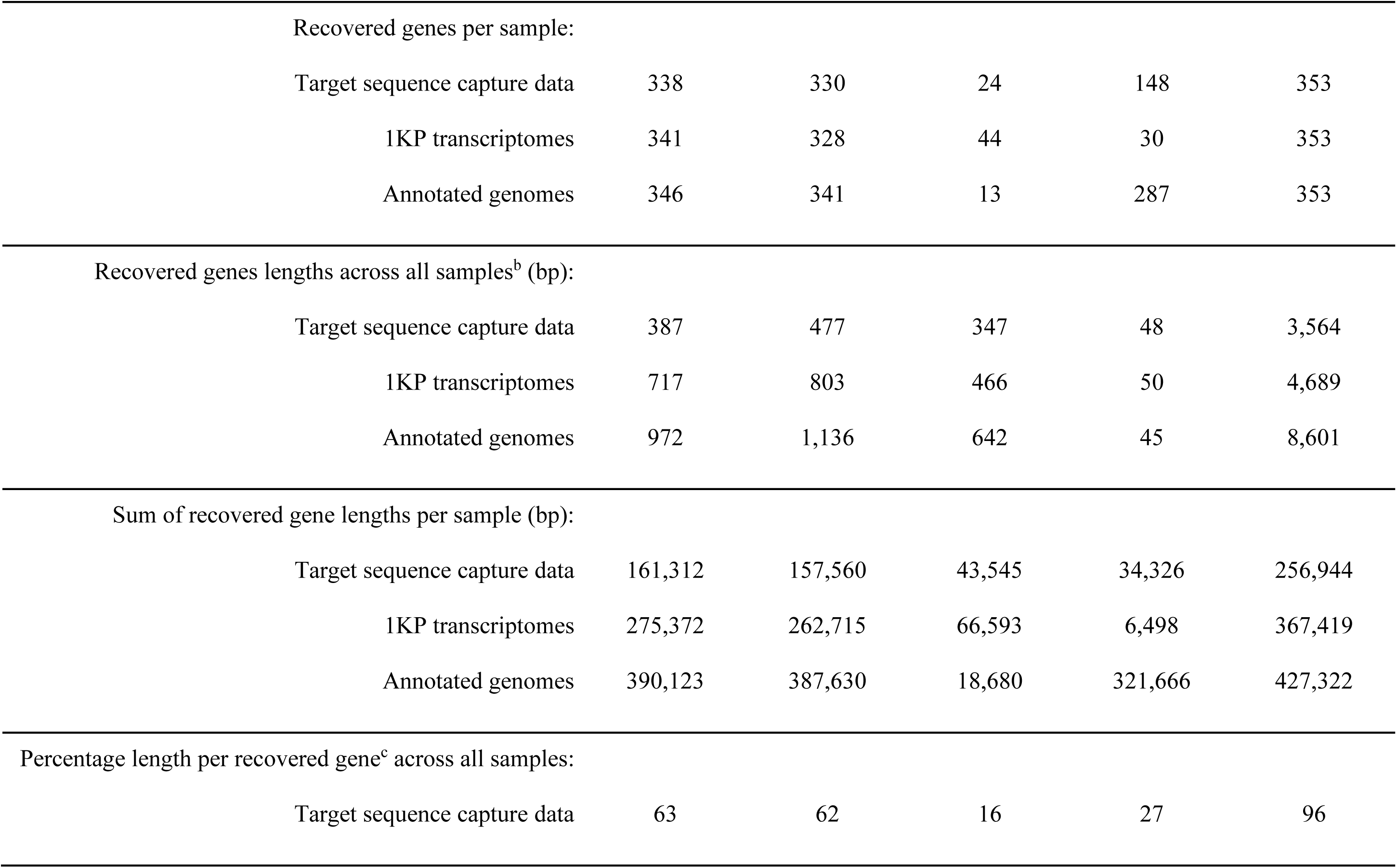
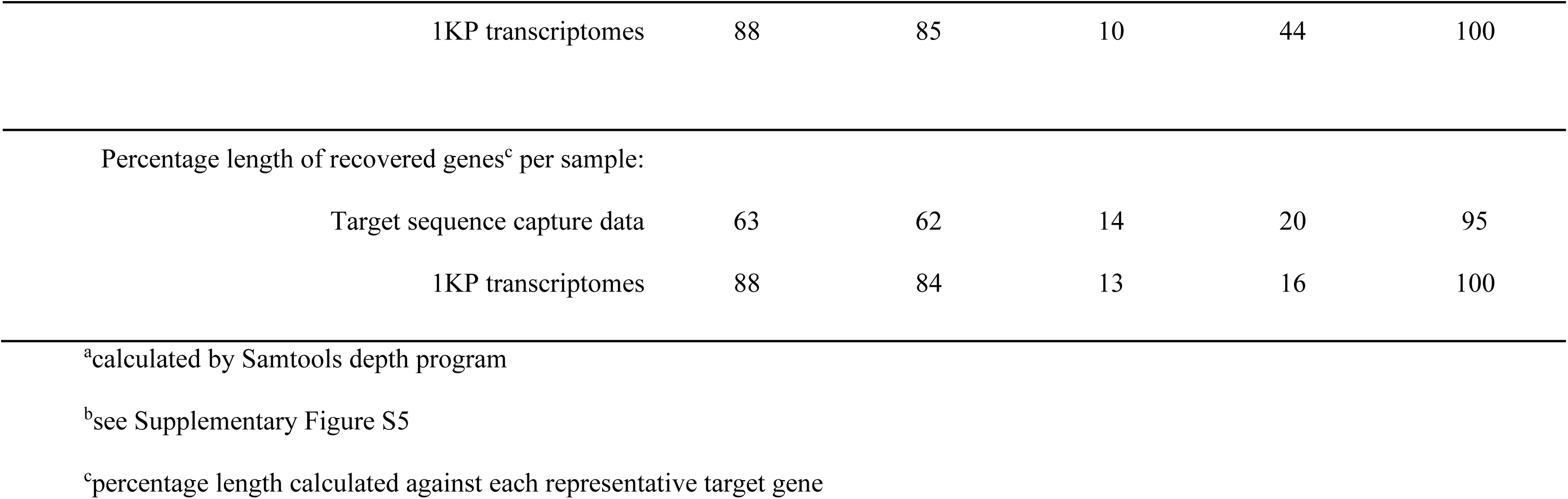
Target sequence capture and gene recovery statistics by sample or gene for Data Release 1.0, including the results of mining of genes from the 1KP and annotated genome datasets. The upper five rows apply to target sequence capture data only.

### Phylogenetic results

The final phylogenetic tree as inferred from Data Release 1.0 is publicly available in interactive form via the Kew Tree of Life Explorer. In the current release, the tree is annotated with local posterior probabilities (LPP, as given by ASTRAL-III) as indicators of branch support. Other measures of support (e.g. quartet scores) can be found within tree files accessible via the RBGK secure FTP. For completeness, the tree is also available in various formats, including Newick (Supplementary Fig. S2).

As a result of filtering and trimming steps during alignment, six genes in Data Release 1.0 were excluded from downstream phylogenetic analysis due to insufficient overlap between sequences. All statistics provided below refer to the remaining dataset. Thus, the species tree is based on 347 gene alignments totalling 824,878 sequences, 489,086,049 base pairs and 532,260 alignment columns. Of these, 509,987 columns (96%) are variable and 475,181 columns (89%) are parsimony informative. The proportion of missing data across all alignments is 61.6% and the median number of genes per sample is 284 (mean: 265.3, standard deviation (SD): 64.3, min: 22, max: 347; Supplementary Table S2). The median number of samples per gene alignment is 2,421 (mean: 2,377.2, SD: 359) and median alignment length is 1,259 (mean: 1,533.9, SD: 985.7; Table 4). The resulting gene trees are highly resolved, with a median support across all nodes (ultrafast bootstrap) of 98% (mean: 87.8%, standard deviation (SD): 18.560) across all nodes in all gene trees (Fig. 7). Only 1.3% of all nodes in all gene trees are very poorly supported (ultrafast bootstrap <30%; Fig. 7) and thus collapsed prior to species tree inference. Further statistics for individual gene alignments and gene trees are reported in Table 4 and Supplementary Table S2.

**Figure 7.**
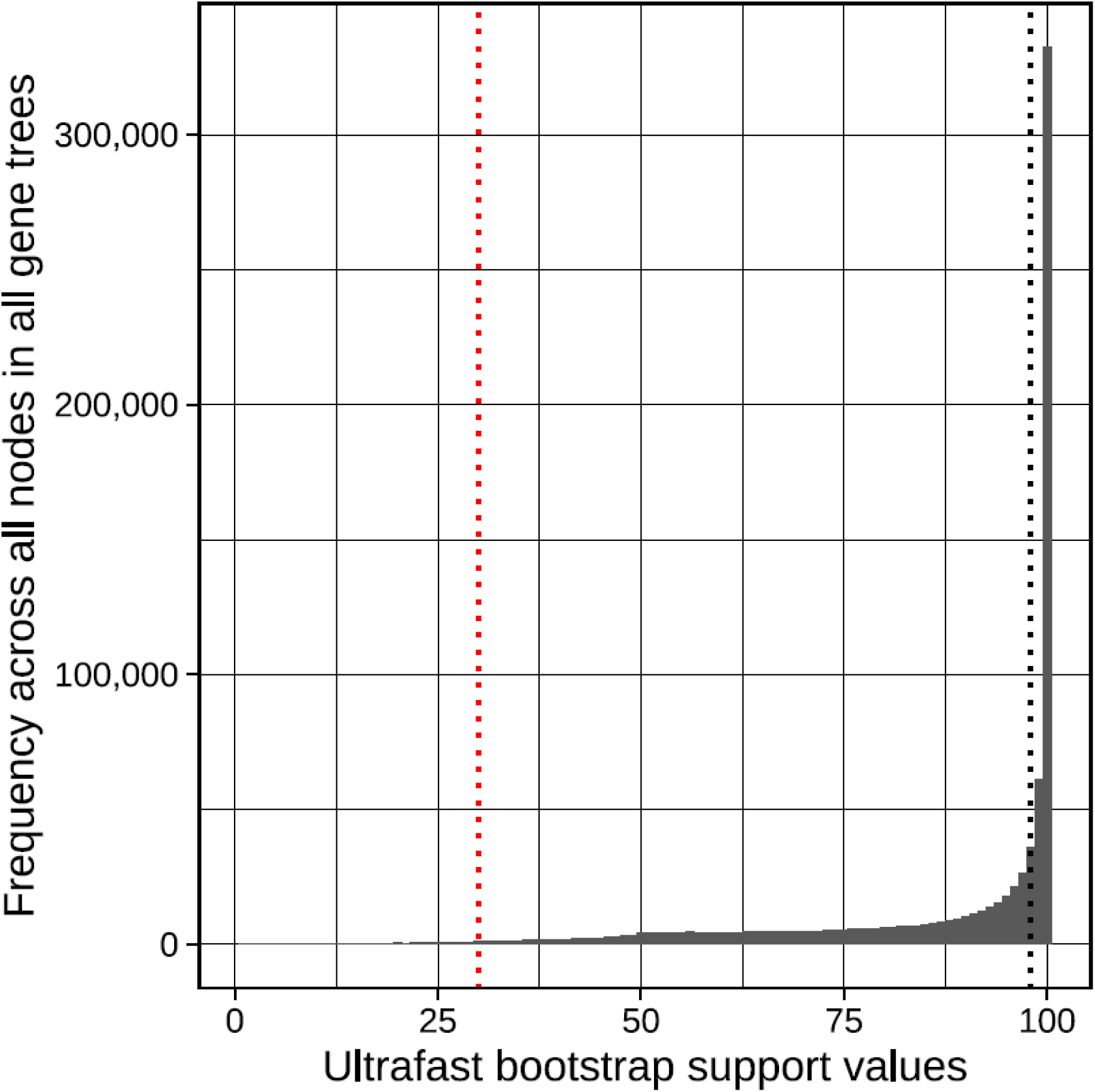
Distribution of ultrafast bootstrap support values across all nodes in all gene trees. Bootstrap values were estimated with IQ-TREE 2.0.5 (Hoang et al. 2017; Minh et al. 2020). Black dotted line indicates the median (98%) and the red dotted line indicates the threshold (30%) for collapsing nodes with low support prior to species tree inference with ASTRAL-III (Zhang et al. 2018). Only 1.3% of all nodes across gene trees are collapsed prior to species tree inference.

**Table 4.** Properties of the 347 gene alignments and gene trees underpinning the species tree included in the Kew Tree of Life Explorer Data Release 1.0.

|  | **Median** | **Mean** | **Standard**<br>**deviation** | **Minimum** | **Maximum** |
| --- | --- | --- | --- | --- | --- |
| Number of samples | 2,421 | 2,377.2 | 358.8 | 491 | 3,014 |
| % of total samples<sup>a</sup> | 77.9 | 76.5 | 11.5 | 15.8 | 96.9 |
| Alignment length | 1,259.0 | 1,533.9 | 985.7 | 250 | 8,119 |
| % missing data<sup>b</sup> | 58.9 | 57.9 | 11.3 | 14.4 | 85.8 |
| Variable sites | 1,224 | 1,469.7 | 940.6 | 240 | 7,873 |
| % variable sites | 96.6 | 96.0 | 2.5 | 81.5 | 100 |
| Parsimony informative sites | 1,137 | 1,369.4 | 859.3 | 233 | 6,792 |
| % parsimony informative<br>sites | 90.7 | 90.0 | 4.20 | 69.1 | 98.9 |
| % nodes in gene trees above<br>30% UFBS<sup>c</sup> | 98.9 | 98.5 | 1.3 | 90.7 | 99.9 |
| Mean support<sup>c</sup> of all nodes | 88.1 | 87.8 | 2.7 | 78.9 | 94.3 |
| Median support<sup>c</sup> of all nodes | 98.0 | 97.6 | 1.8 | 90.0 | 100 |

<sup>a</sup>percentage of samples in species tree present in alignment/gene tree
<sup>b</sup>percentage of empty cells in each alignment
<sup>c</sup>UFBS: ultrafast bootstrap

The species tree accommodates 82% of the quartet relationships in the gene trees (ASTRAL normalized quartet score of 0.82). The majority (76.8%) of nodes in the species tree were well-supported (LPP ≥95%, cf. Sayyari and Mirarab 2016), and only seven nodes were informed by too few gene trees (i.e. <20) to evaluate support. Comparing node support in the species tree at different taxonomic levels (Fig. 8), median quartet support is progressively higher towards shallower taxonomic levels (Fig. 8c), while the effective number of gene trees informing nodes shows the opposite trend (Fig. 8e). Local posterior probabilities show a tendency to be lower (1st quartile) at the deepest taxonomic level (Fig. 8a). Major groups (i.e. monocots, asterids and rosids) show similar distributions of both local posterior probabilities (Fig. 8b) and quartet support values (Fig. 8d), despite the fact that the effective number of gene trees supporting nodes is more variable in monocots (Fig. 8f), which is the result of the lower recovery rates for some orders in this group such as Alismatales, Commelinales and Liliales (Supplementary Fig. S3).

**Figure 8.**
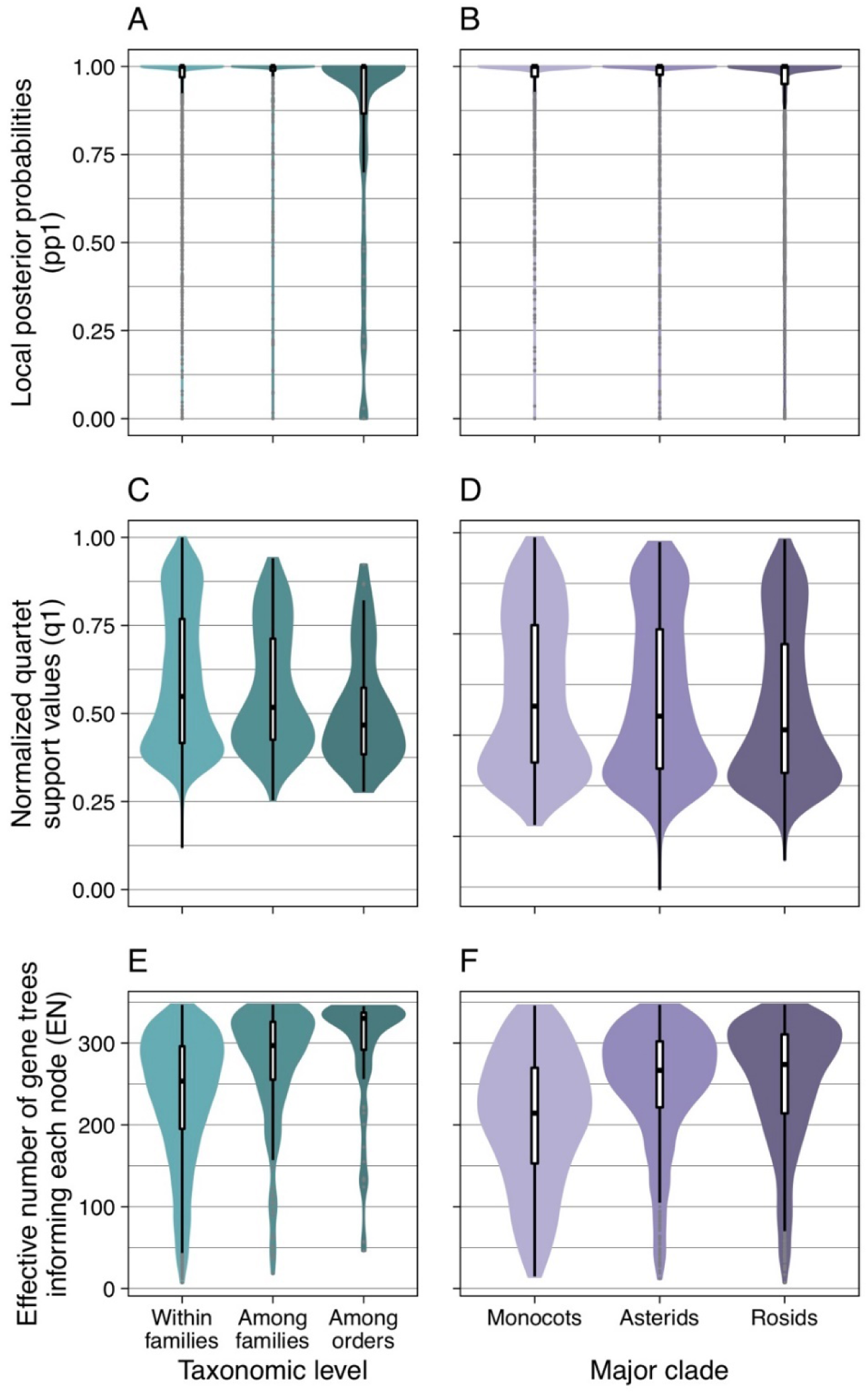
Summary of node properties in the species tree derived from ASTRAL-III (Zhang et al. 2018). Data are grouped by (a, c, e) taxonomic level and (b, d, f) major taxonomic groups. In a, c and e, “within families” refers to relationships within families; “among families” refers to relationships within orders but among families; “among orders” refers to relationships among orders. Box plots show medians, 1^st^ and 3^rd^ quartiles (hinges), and the full distribution excluding outliers (whiskers).

Discounting taxa represented by a single sample (193 families, one order), 96% of testable families and 83% of testable orders were resolved as monophyletic in the species tree. Most of the samples of non-monophyletic families and orders could be assigned to a clade that represents the family or order well, despite lacking some samples and/or containing some outlier samples from other taxa (“concordant taxa” where taxon concordance score >0.5, see Materials and Methods for details). Only five families (Francoaceae, Hernandiaceae, Phyllanthaceae, Pontederiaceae and Schlegeliaceae, represented by 11 samples) and two orders (Bruniales and Icacinales, represented by six samples) were so dispersed that this was not possible (“discordant taxa” where taxon concordance score ≤ 0.5). At the family level, 2,893 samples were resolved in the expected family, two samples were resolved in an unexpected position, and 204 samples were not testable because they belonged to a discordant family or a family represented by a single sample. At the order level, 3,060 samples were resolved in the expected order, 32 samples were resolved in an unexpected position, and seven samples were not testable (see Supplementary Tables S3-S5 for lists of specimens from singly represented taxa, poorly resolved taxa, and outliers to well-resolved taxa, respectively). Placements of all but five genera and seven families were consistent with the WCVP/APG IV taxonomic hierarchy of genera, families and orders. Concordance with existing taxonomy was lower at the genus level, with only 74% of testable genera resolving as monophyletic and 47 genera (represented by 130 samples) being discordant; these numbers partly reflect the deliberate inclusion of multiple samples from genera suspected a priori to be potentially non-monophyletic.

In addition to resolving most genera, families and orders as monophyletic, our tree supports more than half (58%) of the relationships among orders presented by the Angiosperm Phylogeny Group (APG IV 2016; Supplementary Fig. S4). Congruence with APG IV varies among major clades, being notably high in magnoliids (100% of APG IV relationships supported) and monocots (80%), while being substantially lower in eudicots (47%), especially in rosids (33%). Nodes in our tree that are congruent with APG IV ordinal relationships are slightly better supported on average (mean LPP 0.98, median 1) than nodes that are incongruent with APG IV (mean LPP 0.75, median 0.94).

### Tree of Life Explorer

The Kew Tree of Life Explorer (https://treeoflife.kew.org) provides open access to taxon, specimen, sequence, alignment and tree data, with associated metadata for the current data release in accordance with the Toronto guidelines on pre-publication data sharing (Toronto International Data Release Workshop Authors 2009). Users can browse by species, gene or interactive phylogenetic tree. The species interface permits searches by order, family, genus or species, and provides voucher specimen metadata (including links to online specimen images, where available), simple sequence metrics, access to assembled genes and raw data. The gene interface documents all Angiosperms353 genes and associated metrics, links to gene identities in UniProt (https://www.uniprot.org/) and provides access to assembled genes across taxa. The tree of life interface enables browsing and taxon searching of the species tree inferred from the current release dataset, as well as tree downloads (as PNG or Newick) and zooming into user-defined subtrees. All processed data (assembled genes, alignments, gene trees, species trees) and archived releases are available from RBGK’s secure FTP site (http://sftp.kew.org/pub/treeoflife/), whereas raw sequence reads are deposited within the European Nucleotide Archive (project number PRJEB35285) for integration within the Sequence Read Archive.

## DISCUSSION

The new phylogenomic platform described here is a major milestone towards a comprehensive tree of life for all flowering plant species. Firstly, the sequencing of a standardised nuclear marker set of this scale for so many taxa is unprecedented, opening doors to a highly integrated future for angiosperm phylogenetics in the genomic era. Much like a “next generation” *rbcL*, which underpinned so many Sanger sequencing-based plant phylogenetic studies, the Angiosperms353 genes offer opportunities for continuous synthesis of HTS data across angiosperms. The foundational dataset presented here can be re-used or extended for tree of life research at almost any taxonomic scale (Johnson et al. 2019; Larridon et al. 2019; Van Andel et al. 2019; Murphy et al. 2020; Pérez-Escobar et al. 2020; Shee et al. 2020; Slimp et al. 2020; Beck et al. 2021). Secondly, this is the first phylogenetic project to gather novel HTS data across angiosperms with a stratified taxon sampling at the genus level. Our sampling strategy systematically and comprehensively represents both the diversity of angiosperms and their deep-time diversification. As genus-level sampling becomes increasingly complete—a target that is well within reach—this backbone will substantially increase our ability to study the dynamics of plant diversity over time and revisit long-standing questions in systematics (Magallón et al. 2018; Sauquet and Magallón 2018; Soltis et al. 2019). Importantly, it will also sharpen the focus on truly intractable phylogenetic problems (Yang et al. 2020; Zhao et al. 2020), encouraging the exploration of the biological drivers of these phenomena.

Our approach has already led to a burst of community engagement. More than a dozen studies utilising Angiosperms353 probes are already published (e.g. Larridon et al. 2019; Howard et al. 2020; Murphy et al. 2020; Pérez-Escobar et al. 2020; Shee et al. 2020; Slimp et al. 2020; McLay et al. in press), and two journal special issues focused on the probe set are in preparation arising from a recent symposium (Lagomarsino and Jabaily 2020). The probe set has also been adopted by the Genomics for Australian Plants consortium (https://www.genomicsforaustralianplants.com/), which aims to sequence all Australian angiosperm genera, coordinating with the PAFTOL project to optimise collective taxonomic coverage. A subset of the Angiosperms353 genes is now accessible for non-angiosperm land plants thanks to a probe set developed in parallel (Breinholt et al. 2021), inviting the prospect of data integration across all land plants. Angiosperms353 genes (as distinct from the Angiosperms353 probes) are also being leveraged as components of custom-designed probe sets (e.g. Jantzen et al. 2020; Ogutcen et al. 2021). This approach gives all the integrative benefits of Angiosperms353, while permitting (i) the tailoring of Angiosperms353 probes to a specific taxonomic group to increase gene recovery, and (ii) the inclusion of additional loci pertinent to the research in question. Angiosperms353 probes have also been directly combined with an existing custom probe set (Nikolov et al. 2019) as a “probe cocktail” in a single hybridisation, capturing both sets of targets simultaneously with remarkable efficiency (Hendriks et al. in press). These possibilities render the invidious choice between specific and universal probe sets increasingly irrelevant (Kadlec et al. 2017).

We took several open data measures to encourage community uptake, in both the design of our tools and the sharing of our data. The Angiosperms353 probe set itself was designed to be a transparent, “off-the-shelf” toolkit that is open, inexpensive and accessible to all, especially researchers discouraged by the complexity and cost of custom probe design (Johnson et al. 2019). Our sequence data for Angiosperms353 genes are openly available via the Kew Tree of Life Explorer and the Sequence Read Archive, as a public foundation dataset shared according to pre-publication best practice (Toronto International Data Release Workshop Authors 2009). The Explorer offers enhanced transparency and accessibility by allowing users to navigate the data via a phylogenetic snapshot of the current release, along with metadata (e.g. specimen data) and intermediate data (e.g. gene assemblies, alignments, gene trees). Thanks to these resources, cross-community collaboration via Angiosperms353 is gaining momentum.

Our tree, which is based on the most extensive nuclear phylogenomic dataset in flowering plants to date, is strongly supported, credible and highly congruent with existing taxonomy and many hypothesized relationships among orders (APG IV 2016; Supplementary Fig. S4). The data confirm both the effectiveness of Angiosperms353 probes across all major angiosperm clades and the ability of the genes to resolve relationships across taxonomic scales (Fig. 8). Variable sequence recovery notwithstanding (Table 3, Supplementary Fig. S3), most nodes in our tree are underpinned by large numbers of gene trees (Fig. 8e), allowing the species tree to be inferred with confidence (Fig. 8a) despite gene tree conflict (Fig. 8c). However, even the most strongly supported phylogenetic hypotheses must be viewed with caution as they may be biased by model misspecification and wrong assumptions. Moreover, our “first pass” analyses based on a set of standard methods may not suit this dataset perfectly (see below). Nevertheless, our findings are rendered credible by their high concordance with taxonomy, an independent point of reference that has been extensively ground-truthed by pre-phylogenomic DNA data, especially plastid loci.

Agreement with existing family circumscriptions is particularly striking. In contrast, congruence with previously hypothesized relationships among orders (APG IV 2016) is much lower (Supplementary Fig. S4). Some of these earlier hypothesized ordinal relationships derive from relatively weak evidence (bootstrap/jackknife >50%; APG IV 2016), which may partly explain this disagreement. However, it may also be due to phylogenetic conflict between nuclear and plastid genomes, as the established ordinal relationships rest primarily on evidence from plastid loci, substantiated more recently by plastid genomes (Li et al. 2019). It is hardly surprising, then, that a large-scale nuclear analysis presents strongly supported, alternative relationships (Supplementary Fig. S4). The conundrum remains that these incongruences are visible at the ordinal backbone, but not the family level. A more comprehensive exploration of these relationships, the underlying phylogenetic signal and their systematic implications is currently underway.

The analyses presented here are primarily intended as a window onto the information content of our current data release and are not a complete exploration of the data. Thus, downstream application of the current species tree comes with caveats. We used current, widely accepted methods in a pipeline that can be re-run in a semi-automated fashion whenever we release new data. As a consequence, not all possible analysis options and effects have been explored. We anticipate that users of our data will probe it more rigorously and will tailor both sampling and phylogenomic analyses to their specific questions.

Important limitations in our analysis relate to (i) sampling, (ii) gene recovery, (iii) models of sequence evolution and (iv) paralogy. Sampling for intermediate data releases is biased by the current state of progress towards our systematic sampling strategy. This will be addressed in future data releases and can be adjusted by users of our data. Gene recovery relied upon the standard Angiosperms353 target file (Johnson et al. 2019), but it has recently become apparent that tailoring target sequences to taxonomic groups can improve recovery (McLay et al. in press); this will be tested in future releases. Moreover, we are yet to exploit intronic data captured in the “splash zone” adjacent to our target exons. By necessity, our “first pass” phylogenetic analysis does not explore the fast-evolving spectrum of methodological options available for phylogenomic analysis. For example, we rely on a simple standard model of sequence evolution, but more sophisticated models accounting for codon positions or amino acids may improve phylogenetic inference. Potential paralogy is not addressed by our current pipeline. The genes underpinning our analysis were carefully chosen to represent single-copy genes across flowering plants (Johnson et al. 2019; Leebens-Mack et al. 2019). However, some paralogy may have gone unnoticed due to the pervasiveness of gene and genome duplication in plants (Li and Barker 2020). Overall, we expect that the occasional presence of paralogs in our current analysis would more likely lead to inflated estimates of gene tree incongruence, and thus result in reduced support values, than significant topological biases (Yan et al. 2020). Thus, we consider our tree relatively conservative while acknowledging that we are not yet exploiting the full potential of our data. Although a rigorous analysis of paralogy in Angiosperms353 genes was not tractable for this data release, we look forward to deeper insights emerging as community-wide engagement with Angiosperms353 grows.

## PROSPECTS

In the immediate future, we will deliver a further data release through which we expect to reach the milestone of sampling 50% of all angiosperm genera. This target will be achieved through substantial novel data production by PAFTOL and collaborators, augmented by data mined from public sources. In-depth phylogenetic analyses of our data and their evolutionary implications are also underway.

Beyond this point, we see three priority areas in which future platform developments might be concentrated, resources permitting. Firstly, taxon sampling to the genus level must be completed. Our original target of sampling all angiosperm genera remains, but the mode of reaching this is likely to evolve. We anticipate an acceleration in production of Angiosperms353 data by the broader community. The completion of generic-level sampling will require both the integration of community data in the broader angiosperm tree of life as well as strategic investment in filling inevitable data gaps for orphan groups. Secondly, numerous opportunities for refinement exist across our methods. For example, insights from our data might permit the optimisation of the Angiosperms353 probes to improve gene capture. Efficiency of gene assembly from sequence data can also be improved bioinformatically (McLay et al. in press). As costs of sequencing decline, target sequence capture *in vitro* may no longer be necessary, the target genes being retrieved simply from sufficiently deeply sequenced genomes. Thirdly, for the full integrative potential of Angiosperms353 genes to be achieved, infrastructure for aggregating and sharing this coherent body of data must be improved. While the Kew Tree of Life Explorer provides a proof-of-concept, it is the public data repositories (e.g. NCBI, ENA) that offer the greatest prospects of a mechanism to achieve this. To fully parallel the earlier success of public repositories for facilitating single-gene phylogenetic trees (e.g. *rbcL*, *matK*), new tools are needed to assist with efficient upload and annotation of target capture loci and associated metadata.

Even with a completed genus-level angiosperm tree of life well within reach, the monumental task of sampling all species remains. The scale of this challenge is 24-fold greater than the genus-level tree towards which we are currently working. However, with sufficient investment, increased efficiencies and community engagement, such an ambition could potentially be realised. Collections-based institutions are poised to play a critical role in this endeavour through increasingly routine molecular characterisation of their specimens, perhaps as part of digitisation programmes, and are already facilitating the growing trend towards species-complete sampling in phylogenomic studies (e.g. Loiseau et al. 2019; Murphy et al. 2020; Kuhnhäuser et al. 2021). Our platform demonstrates how large-scale phylogenomic projects can capitalise on natural history collections to achieve a much more complete sampling than hitherto possible.

The growing movement to sequence the genomes of all life on Earth, inspired by the Earth Biogenome Project (Lewin et al. 2018), significantly boosts the prospects for completing the tree of life for all species, but is hampered by the focus on “gold standard” whole genomes requiring the highest quality input DNA. Our platform offers the opportunity to bridge the gap between the ambition of these projects and the vast phylogenomic potential of natural history collections. However, as life on Earth becomes increasingly imperilled, we cannot afford to wait. To meet the urgent demand for best estimates of the tree of life, we must dynamically integrate phylogenetic information as it is generated, providing synthetic trees of life to the broadest community of potential users (Eiserhardt et al. 2018). Our platform facilitates this crucial synthesis by providing a cross-cutting dataset and directing the community towards universal markers that seem set to play a central role in completing an integrated angiosperm tree of life.

## DATA AVAILABILITY AND SUPPLEMENTARY MATERIAL

All data generated in this study are publicly released under a Creative Commons Attribution 4.0 International (CC BY 4.0) license and the Toronto guidelines on pre-publication data sharing (Toronto International Data Release Workshop Authors 2009). The data are accessible via the Kew Tree of Life Explorer (https://treeoflife.kew.org) and our secure FTP (http://sftp.kew.org/pub/treeoflife/). Raw sequence reads are deposited in the European Nucleotide Archive (https://www.ebi.ac.uk/ena/browser/home) under umbrella project PRJEB35285. Scripts and other files relating to our phylogenomic pipeline are available at our GitHub (https://github.com/RBGKew/KewTreeOfLife). Supplementary materials cited in this paper are available from the Dryad Digital Repository (http://dx.doi.org/10.5061/dryad.[NNNN]).

## FUNDING

This work was supported by grants from the Calleva Foundation and the Sackler Trust to the Plant and Fungal Trees of Life project at the Royal Botanic Gardens, Kew. Additional support was received from the Garfield Weston Foundation, as part of the Global Tree Seed Bank Programme.

## ACKNOWLEDGEMENTS

We would like to thank Guilherme Antar, Alex Antonelli, Marc Appelhans, Julien Bachelier, Donovan Bailey, Aurélien Bour, Peter Boyce, Gemma Bramley, Sven Buerki, Stuart Cable, Martin Callmander, Monica Carlsen, Vinicius Castro Sousa, Mark Chase, Martin Cheek, Maarten Christenhusz, Thomas Couvreur, Darren Crayn, Iain Darbyshire, Alison Devault, Manuel de la Estrella, Elton John de Lirio, Jurriaan de Vos, Zacky Ezedin, Federico Fabriani, Mike Fay, Geneviève Ferry, Helen Fortune-Hopkins, Jocelyn Hall, Ameka Gabriel Komla, Jim Leebens-Mack, Elliot Gardner, Ester Gaya, Mark Gibernau, Olwen Grace, Sean Graham, Jan Hackel, Anna Haigh, Kasper Hendriks, Oriane Hidalgo, Elizabeth Joyce, Bente Klitgaard, Sophie Lane, Isabel Larridon, Drew Larson, Frederic Lens, Christine Leon, Gwil Lewis, Jing-Xia Liu, Meng Lu, Jaquelini Luber, Eve Lucas, Penny Malakasi, Vidal Mansano, Laura Martinez-Suz, Angela McDonell, Alexander Monro, Michael Moore, Klaus Mummenhof, Tuula Niskanen, Andres Orejuela, Luis Palazzesi, Joe Parker, Frederic Pautz, Jaume Pellicer, Oscar Perez Escobar, Yohan Pillon, Jose Pirani, Robyn Powell, Natalia Przelomska, Carmen Puglisi, Eric Roalson, Hervé Sauquet, Hanno Schaefer, Ruud Scharn, Rowan Schley, David Scherberich, Toral Shah, Mark P. Simmons, Ana Rita Simões, Lalita Simpson, Stephen Smith, Doug Soltis, Pam Soltis, Cynthia Sothers, Marybel Soto Gomez, Jemma Taylor, Liam Trethowan, Anna Trias-Blasi, Tim Utteridge, Juan Viruel, Maria Vorontsova, Gane Ka-Shu Wong, Sin Yeng Wong and Sue Zmarzty for helping PAFTOL reach its goals through collaboration, sharing expertise and providing samples; Noelia Alvarez de Roman, Richard Barley, Nicola Biggs, Elisa Biondi, Elinor Breman, Hannah Button, Christopher Cockel, David Cooke, Nina Davies, Solene Dequiret, John Dickie, Florence Ducan-Antoine, Sara Edwards, Thomas Freeth, Sue Frisby, Tim Fulcher, Aurélie Grall, Anthony Hall, Alex Hankey, Kate Hardwick, Keegan Hickey, David Hickmott, Rebecca Hilgenhof, Imalka Kahandawala, Lara Jewitt, Laura Jennings, Nick Johnson, Udayangani Liu, Carlos Magdalena, Max Moog, Richard Moore, Ana Oliveira, Tim Pearce, Tom Pickering, Sara Redstone, Greg Redwood, Luxy Reed, Paul Rees, Matthew Rees, Silke Roch, Daniel Rosenberg, Marcello Sellaro, Scott Taylor, Janet Terry, Michael Way, Ian Willey, Patricia Woods, Rosie Woods and Martin Xanthos for support with acquisition samples from RBGK collections, both living and preserved; Alexander Bowles, Dion Devey, Laszlo Csiba, Isabel Fairlie, Lorna Frankel, Karime Gutierrez, Alina Höwener, Izai A. B. Sabino Kikuchi, Beata Klejevskaja, Jake Newitt, Michelle Siros and Jessica Tengvall, Haydn Thompson, for assistance with laboratory work and data collection; Laura Green, Alan Paton, Sarah Phillips and Marie-Helene Weech for support with specimen digitisation; Nicholas Black, Michael Bradford, Carol Sinker, Robert Turner and Noor Al Wattar for assistance with computational infrastructure. Finally, special thanks to Kathy Willis, former Director of Science at RBGK, for inspiring the establishment of the PAFTOL project.

## Notes

https://treeoflife.kew.org

http://sftp.kew.org/pub/treeoflife/

https://github.com/RBGKew/KewTreeOfLife

https://www.ebi.ac.uk/ena/browser/view/PRJEB35285

### Competing Interest Statement

The authors have declared no competing interest.

