## Supplementary Materials for "A Comprehensive Phylogenomic Platform for Exploring the Angiosperm Tree of Life"

### Supplementary Methods for Family Identification Validation by DNA barcoding

Novel target sequence capture data and the One Thousand Plant Transcriptomes Initiative (1KP) (Leebens-Mack et al. 2019) samples were validated at family level through in-silico DNA barcoding, using a pipeline that is published on our [Github](https://github.com/RBGKew/KewTreeOfLife).

##### **Samples**

For target sequence capture data, plastomes and ribosomal DNA were recovered from raw reads using GetOrganelle (Jin et al. 2020). In both cases, recommended parameters were used (see https://github.com/Kinggerm/GetOrganelle#recipes; i.e. -R 20 -k 21,45,65,85,105 for plastomes, and -R 10 -k 35,85,115 for nuclear ribosomal DNA).

Samples from 1KP were also validated by barcoding. For these 1KP data, transcriptome assemblies were directly used in barcode tests.

##### **DNA barcode reference databases**

Due to the partial recovery of organellar DNA and the uneven coverage of reference barcodes across families, six barcode tests were performed against individual barcode reference databases that were built from the NCBI Nucleotide database ([https://www.ncbi.nlm.nih.gov](https://www.ncbi.nlm.nih.gov/nuccore)/nuccore) and BOLD (https://www.boldsystems.org), five for specific loci (nuclear ribosomal 18S, as well as plastid *rbcL, matK, trnL*, and *trnH-psbA*) and the remaining one for whole plastomes ([https://ftp.ncbi.nlm.nih.gov/refseq/
release/plastid/](https://ftp.ncbi.nlm.nih.gov/refseq/release/plastid/)).

BOLD was the primary choice for building reference barcode databases as it is curated (Ratnasingham and Hebert 2007). However, only four barcode loci had sufficient coverage across angiosperms (*matk, rbcL, rbcLa* and ITS2), and initial barcode tests with *rbcL* and ITS2 were unsatisfactory, resulting in just two barcode reference databases sourced from BOLD with the other four from NCBI (Table A). NCBI search queries were downloaded using Entrez-direct (Kans 2010) and are available on our [GitHub](https://github.com/RBGKew/KewTreeOfLife). The *trnL* and *trnH-psbA* sequences were trimmed in Cutadapt (Martin 2011) using universal barcode primers with 10% mismatch tolerated. For *trnH-psbA*, primers *psbAf* and *trnH2* were used (Sang et al. 1997; Tate and Simpson 2003), and for *trnL*, primers *trnL-c* and *trnL*-*d* were used (Taberlet et al. 2007). All sequences were further filtered based on length, with a minimum length set for each barcode as in Table A.

| Source | Barcode | Min. length (bases) | Min. coverage  (%) |
| --- | --- | --- | --- |
| NCBI | 18S | 1000 | 90 |
| NCBI | Plastome | 1000 | 0 |
| NCBI | *trnH-psbA* | 200 | 90 |
| NCBI | *trnL* | 200 | 90 |
| BOLD | *matK* | 300 | 90 |
| BOLD | *rbcLa* | 300 | 90 |

**Table A*.*** *Details of the six barcode reference databases used for family identification. BLASTN results were further filtered based on minimum (min.) length and minimum coverage of subject sequence.*

##### **Taxonomic standardization**

The species names associated with the target sequence capture and 1KP samples, and with the sequences from the barcode reference databases, were standardized against the World Checklist of Vascular Plants (WCVP, <https://wcvp.science.kew.org/>) using a custom python script available on [Github](https://github.com/RBGKew/KewTreeOfLife). Sequences from the barcode reference databases whose taxonomic assignment could not be unambiguously resolved to the genus level were discarded.

##### **Barcode tests**

Sample sequences (i.e. plastomes and ribosomal DNA for target sequence capture data, transcriptomes for 1KP data) were queried against the barcode databases using BLASTN (Camacho et al. 2009), only if their family was present in the database. BLASTN results were filtered requiring a minimum identity of >95% and a minimum coverage of reference locus (see Table A). However, BLASTN results against the plastome database were filtered on alignment length, requiring 1,000 bases (note that the median length of plastome contigs recovered from the target sequence capture data was 1,327 bases).

For each locus where the validation could be performed, a sample was considered to have passed the test if the BLASTN match with the highest sequence identity came from the same family as the sample and failed otherwise. The family assignment of the sample was then considered as: (i) Confirmed: if one or more barcode tests matched the family identification of a sample; (ii) Rejected: if more than half of the barcode tests gave the same incorrect family identification (requires at least two barcode tests); (iii) Inconclusive (otherwise).

### Supplementary Results for Family Identification Validation by DNA Barcoding

##### **Target sequence capture data validation by DNA barcoding**

Plastome and ribosomal DNA sequences were recovered for 2,394 and 2,349 out of 2,438 samples, respectively. For ribosomal DNA, the median sum of contigs length was 6.789 kb (expecting ~8 kb), distributed in no more than two contigs 86% of the time (range:1-4 contigs). For plastomes, recovery was less comprehensive with a median sum of contigs length of 32.284 kb (expecting ~150 kb) and fragmented in 21 contigs on average (range: 1-68 contigs).

The presence of the genus and family in the target sequence capture sample were checked in each barcode reference database. On average, the named family for 69% of samples was represented, but the named genus was only represented for 35% of samples. These results varied widely by locus, with *matK* having the best coverage of the named taxa, and *trnH-psbA* and whole plastome sequences having the lowest coverage (Table B).

| **Barcode** | **Genus (%)** | **Family (%)** |
| --- | --- | --- |
| *rbcLa* | 28.7 | 57.7 |
| *matK* | 64.2 | 93.6 |
| *trnL* | 54.6 | 71.8 |
| *trnH-psbA* | 19.1 | 48.7 |
| 18S | 23.4 | 85.2 |
| Plastome | 19.1 | 59.4 |

**Table B.** *Percentage of genera and families in the target sequence capture samples represented in each barcode database.*

Using the validation criteria detailed above, 1,977 out of 2,438 samples were validated (81%) and 19 rejected (0.8%) at family level. A substantial proportion of samples (444, 18%) could not be validated either because their named families were not represented in the barcode databases, or because no BLASTN matches were obtained, at any of the tested loci.

Tests against whole plastomes and 18S ribosomal DNA were conducted most frequently but these had a higher rate of failure than the other tests. In most of these failures, a BLASTN match to a sequence from the expected family was obtained, but the match with highest identity was to a sequence from another family.

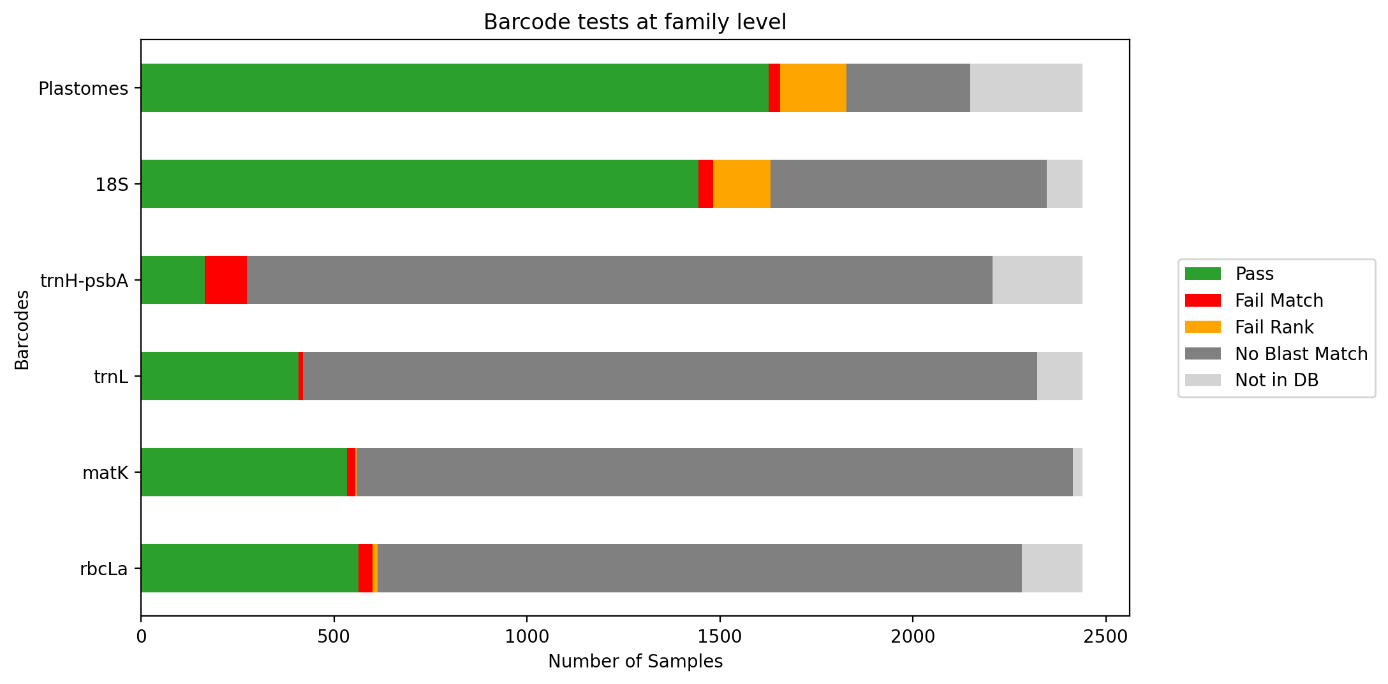

**Figure A**. Results for each barcode test at family level for the target sequence capture samples. The figure shows the number of samples that passed a test (Pass), failed because their family was not in the BLASTN matches (Fail match), failed because their family was in the BLASTN matches but did not rank first (Fail rank), had no BLASTN matches after filtering (No BLASTN match), or did not have their family in the barcode database (Not in DB).

##### **1KP data validation by DNA barcoding**

The validation by barcoding of 1KP samples confirmed 663 out of the 678 samples at family level, rejecting 10 with the remaining five found to be inconclusive (Table C). Note that all 10 rejected samples were excluded from the tree after family identification validation.

| **Sample** | **Barcode validation** | **Family** | **Species** | **Number of barcode tests** | **Matching family in barcode tests** |
| --- | --- | --- | --- | --- | --- |
| FONV | Rejected | Francoaceae | *Greyia sutherlandii* | 4 | Menispermaceae |
| JBGU | Rejected | Amaranthaceae | *Amaranthus palmeri* | 5 | Poaceae |
| KTWL | Rejected | Montiniaceae | *Kaliphora madagascariensis* | 3 | Rubiaceae |
| LVNW | Rejected | Menispermaceae | *Cocculus laurifolius* | 4 | Phyllanthaceae |
| OKEF | Rejected | Dilleniaceae | *Hibbertia grossulariifolia* | 2 | Chloranthaceae |
| OTAN | Rejected | Hydrangeaceae | *Deutzia scabra* | 4 | Gunneraceae |
| PGKL | Rejected | Rutaceae | *Phellodendron amurense* | 6 | Juglandaceae |
| QJXB | Rejected | Thymelaeaceae | *Wikstroemia indica* | 6 | Pittosporaceae |
| RTTY | Rejected | Salvadoraceae | *Salvadora sp.* | 5 | Capparaceae |
| ULGV | Rejected | Rubiaceae | *Morinda citrifolia* | 6 | Annonaceae |
| CYVA | Inconclusive | Ranunculaceae | *Actaea racemosa* | 3 |  |
| PVGM | Inconclusive | Oncothecaceae | *Oncotheca balansae* | 0 |  |
| RDYY | Inconclusive | Tecophilaeaceae | *Cyanastrum cordifolium* | 0 |  |
| SART | Inconclusive | Xeronemataceae | *Xeronema callistemon* | 0 |  |
| YZGX | Inconclusive | Cyrillaceae | *Cyrilla racemiflora* | 2 |  |

**Table C*.*** *Rejected and inconclusive samples, including the number of barcode tests conducted for each sample, and the families they matched best in these tests.*

The two main differences with target sequence capture samples (see above) were (i) that barcoding loci were more frequently recoverable from transcriptome data than from the target sequence capture data, and (ii) that more of the genera tested were present in reference databases (84% of samples at family level, 70% at genus level). See Table D below.

| **Barcode** | **Genus (%)** | **Family (%)** |
| --- | --- | --- |
| *rbcLa* | 68.3 | 78.8 |
| *matK* | 94.9 | 98.8 |
| *trnL* | 84.8 | 88.0 |
| *trnH-psbA* | 51.8 | 72.0 |
| 18S | 60.0 | 92.6 |
| Plastome | 58.0 | 75.6 |

**Table D*.*** *Percentage of genera and families in the 1KP samples represented in each barcode database*

Most barcode tests could be conducted on most samples, with the exception of the *trnH-psbA* locus, for which few BLASTN matches were found (Fig. B).

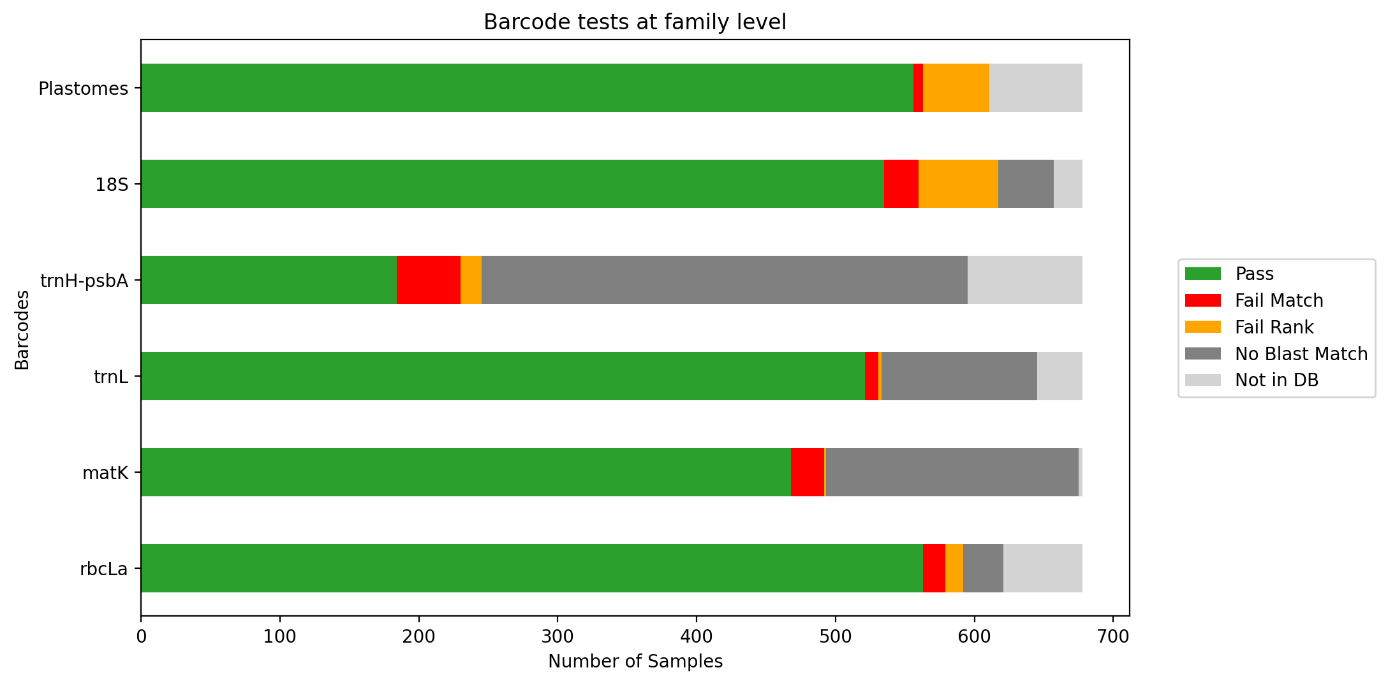

**Figure B**. Results for each barcode test for the 1KP samples. The figure shows the number of samples that passed a test (Pass), failed because their family was not in the BLASTN matches (Fail match), failed because their family was in the BLASTN matches but did not rank first (Fail rank), had no BLASTN matches after filtering (No BLASTN match), or did not have their family in the barcode database (Not in DB).

Leebens-Mack J.H., Barker M.S., Carpenter E.J., Deyholos M.K., Gitzendanner M.A., Graham S.W., Grosse I., Li Z., Melkonian M., Mirarab S., Porsch M., Quint M., Rensing S.A., Soltis D.E., Soltis P.S., Stevenson D.W., Ullrich K.K., Wickett N.J., DeGironimo L., Edger P.P., Jordon-Thaden I.E., Joya S., Liu T., Melkonian B., Miles N.W., Pokorny L., Quigley C., Thomas P., Villarreal J.C., Augustin M.M., Barrett M.D., Baucom R.S., Beerling D.J., Benstein R.M., Biffin E., Brockington S.F., Burge D.O., Burris J.N., Burris K.P., Burtet-Sarramegna V., Caicedo A.L., Cannon S.B., Çebi Z., Chang Y., Chater C., Cheeseman J.M., Chen T., Clarke N.D., Clayton H., Covshoff S., Crandall-Stotler B.J., Cross H., DePamphilis C.W., Der J.P., Determann R., Dickson R.C., Di Stilio V.S., Ellis S., Fast E., Feja N., Field K.J., Filatov D.A., Finnegan P.M., Floyd S.K., Fogliani B., García N., Gâteblé G., Godden G.T., Goh F. (Qi Y., Greiner S., Harkess A., Heaney J.M., Helliwell K.E., Heyduk K., Hibberd J.M., Hodel R.G.J., Hollingsworth P.M., Johnson M.T.J., Jost R., Joyce B., Kapralov M. V., Kazamia E., Kellogg E.A., Koch M.A., Von Konrat M., Könyves K., Kutchan T.M., Lam V., Larsson A., Leitch A.R., Lentz R., Li F.W., Lowe A.J., Ludwig M., Manos P.S., Mavrodiev E., McCormick M.K., McKain M., McLellan T., McNeal J.R., Miller R.E., Nelson M.N., Peng Y., Ralph P., Real D., Riggins C.W., Ruhsam M., Sage R.F., Sakai A.K., Scascitella M., Schilling E.E., Schlösser E.M., Sederoff H., Servick S., Sessa E.B., Shaw A.J., Shaw S.W., Sigel E.M., Skema C., Smith A.G., Smithson A., Stewart C.N., Stinchcombe J.R., Szövényi P., Tate J.A., Tiebel H., Trapnell D., Villegente M., Wang C.N., Weller S.G., Wenzel M., Weststrand S., Westwood J.H., Whigham D.F., Wu S., Wulff A.S., Yang Y., Zhu D., Zhuang C., Zuidof J., Chase M.W., Pires J.C., Rothfels C.J., Yu J., Chen C., Chen L., Cheng S., Li J., Li R., Li X., Lu H., Ou Y., Sun X., Tan X., Tang J., Tian Z., Wang F., Wang J., Wei X., Xu X., Yan Z., Yang F., Zhong X., Zhou F., Zhu Y., Zhang Y., Ayyampalayam S., Barkman T.J., Nguyen N. phuong, Matasci N., Nelson D.R., Sayyari E., Wafula E.K., Walls R.L., Warnow T., An H., Arrigo N., Baniaga A.E., Galuska S., Jorgensen S.A., Kidder T.I., Kong H., Lu-Irving P., Marx H.E., Qi X., Reardon C.R., Sutherland B.L., Tiley G.P., Welles S.R., Yu R., Zhan S., Gramzow L., Theißen G., Wong G.K.S. 2019. One thousand plant transcriptomes and the phylogenomics of green plants. Nature. 574:679–685.

Martin M. 2011. Cutadapt removes adapter sequences from high-throughput sequencing reads. EMBnet.journal. 17:10–12.

Ratnasingham S., Hebert P. 2007. The Barcode of Life Data System (www.barcodinglife.org). Mol. Ecol. Notes. 7:355–364.

Sang T., Crawford D.J., Stuessy T.F. 1997. Chloroplast DNA phylogeny, reticulate evolution, and biogeography of *Paeonia* (Paeoniaceae). Am. J. Bot. 84:1120-1136.

Taberlet P., Coissac E., Pompanon F., Gielly L., Miquel C., Valentini A., Vermat T., Corthier G., Brochmann C., Willerslev E. 2007. Power and limitations of the chloroplast trnL (UAA) intron for plant DNA barcoding. Nucleic Acids Res. 35:e14.

Tate J.A., Simpson B.B. 2003. Paraphyly of *Tarasa* (Malvaceae) and diverse origins of the polyploid species. Syst. Bot. 28:723-737.

### Supplementary Figures

#### **Supplementary Figure S1**

Illustration of the phylogenetic validation. Taxon concordance scores are calculated to identify the node best representing each family in the tree (or order if the family is represented by a single sample). In this example, samples A1 to A3 belong to family A and B1 to B3 to family B.  In step 1, concordance scores are calculated (here illustrated for family A). The node marked with a green dot is the highest scoring for this family and is thus its best representative, although the taxon does not resolve as a monophyletic group. In Step 2, samples A1 to A3 are found under that node and are thus confirmed.  The node marked with a blue dot best represents family B and samples B2 and B3 are thus confirmed. However, sample B1 is not found under that node and is thus rejected.

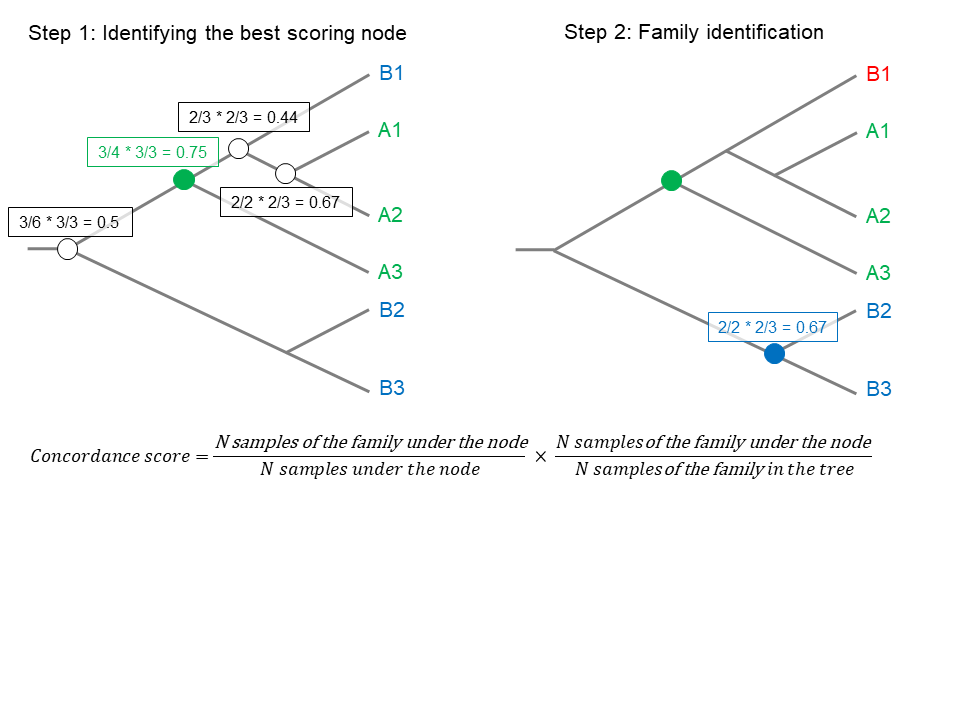

#### **Supplementary Figure S2**

Flowering plant phylogenetic tree. Each tip is labelled with order, family, genus, species and sample ID (ENA Run Accession for target sequence capture data, NCBI Project Accession for Annotated Genomes, Sample ID for 1KP). Branches are labelled with local posterior probability support values.

This figure is available in three formats for visualizing the tree: (i) Supplementary Fig. S2.html, (ii) Supplementary Fig. S2.svg and (iii) Supplementary Fig. S2.pdf. The figure can also be downloaded as a nexus file: Supplementary Fig. S2.nex

#### **Supplementary Figure S3**

Recovery summaries per sample for all target sequence capture data grouped by order and superimposed on a simplified tree showing phylogenetic relationships between orders (a). The plots show the distribution of (b) the sum of gene lengths in base pairs, (c) the number of recovered genes, and (d) the number of retained genes in final alignment. Values for orders with fewer than three samples are shown as points.

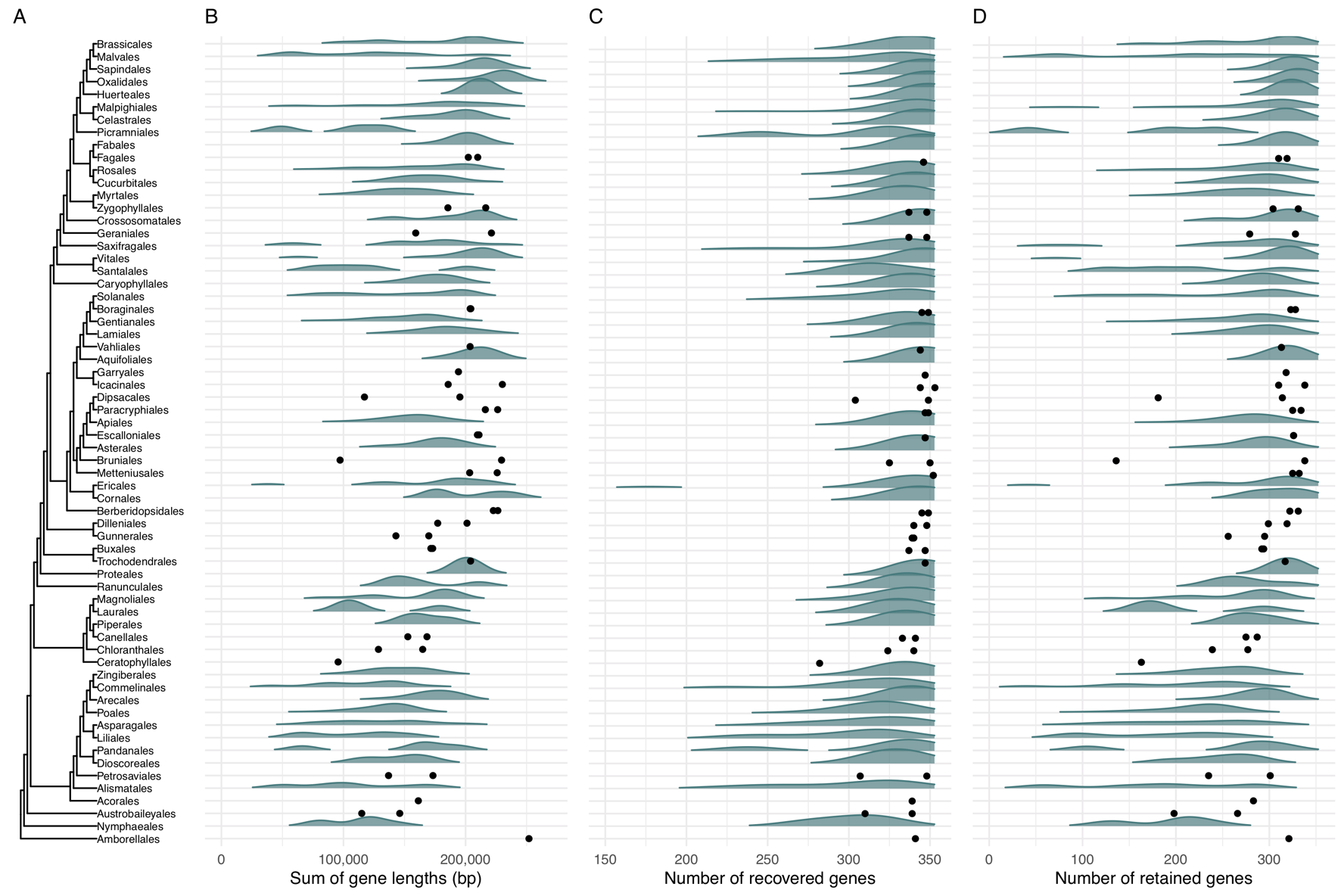

#### **Supplementary Figure S4**

Tanglegram comparing the Angiosperms353 and APG IV phylogenetic trees at ordinal level. Nodes in the Angiosperms353 tree are labelled with support values (local posterior probability, LPP); unlabelled nodes are maximally supported (LPP=1). Nodes in the APG IV tree are labelled with blue dots where they are congruent with nodes in the Angiosperms353 tree, and red dots if they are in conflict. For orders that are not monophyletic in the Angiosperms353 tree, congruence with the APG IV is calculated based on their most compatible placement in the Angiosperms353 tree, disregarding all other placements; thus, a node is treated as congruent whenever possible.

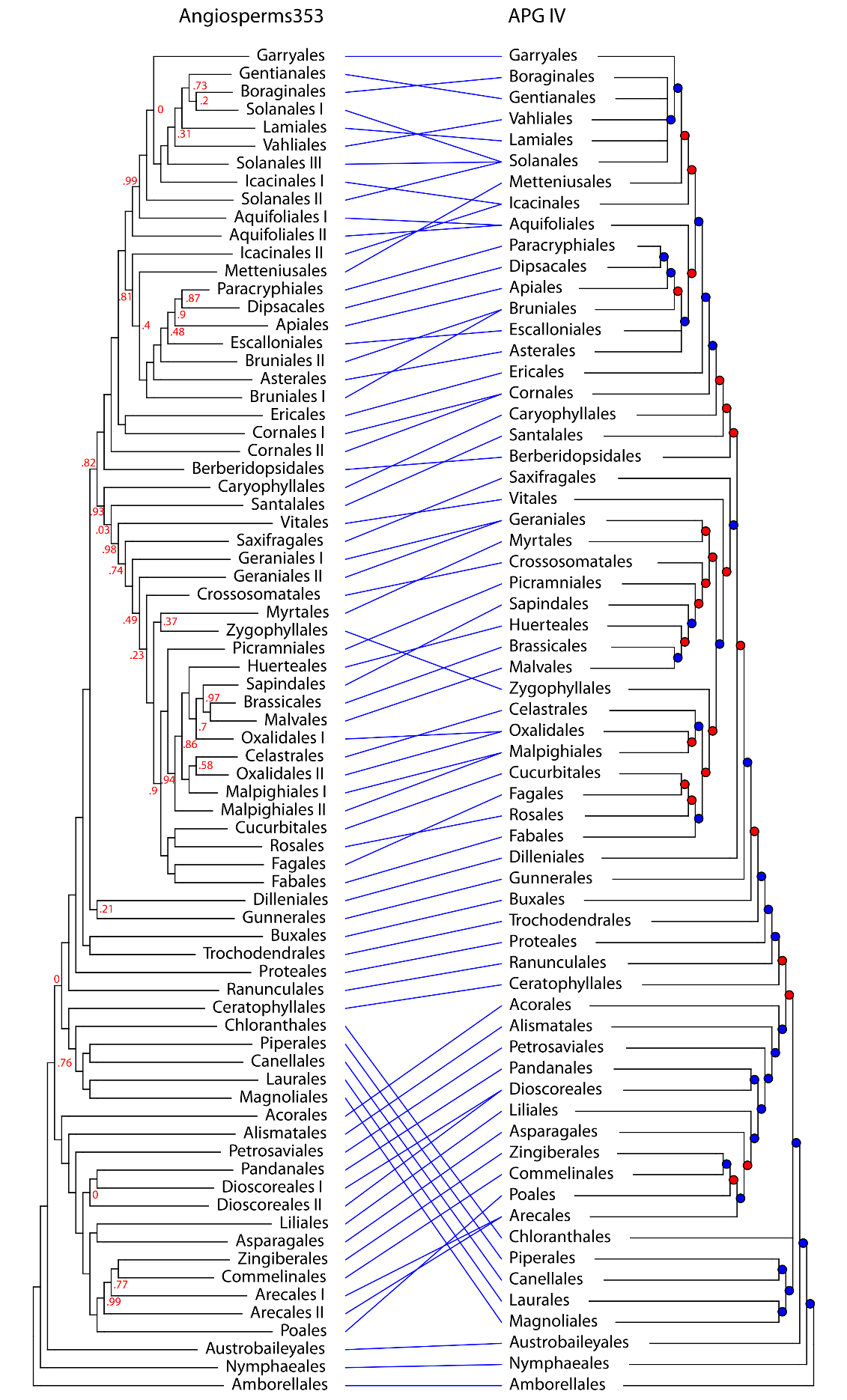

#### **Supplementary Figure S5**

Distribution of recovered gene lengths across all samples in Data Release 1.0, grouped by data source. The black dotted line indicates overall median (461, mean: 560.4, SD: 414.8, min: 45, max: 8601). For detailed statistics per data source, see Table 3 in main text.

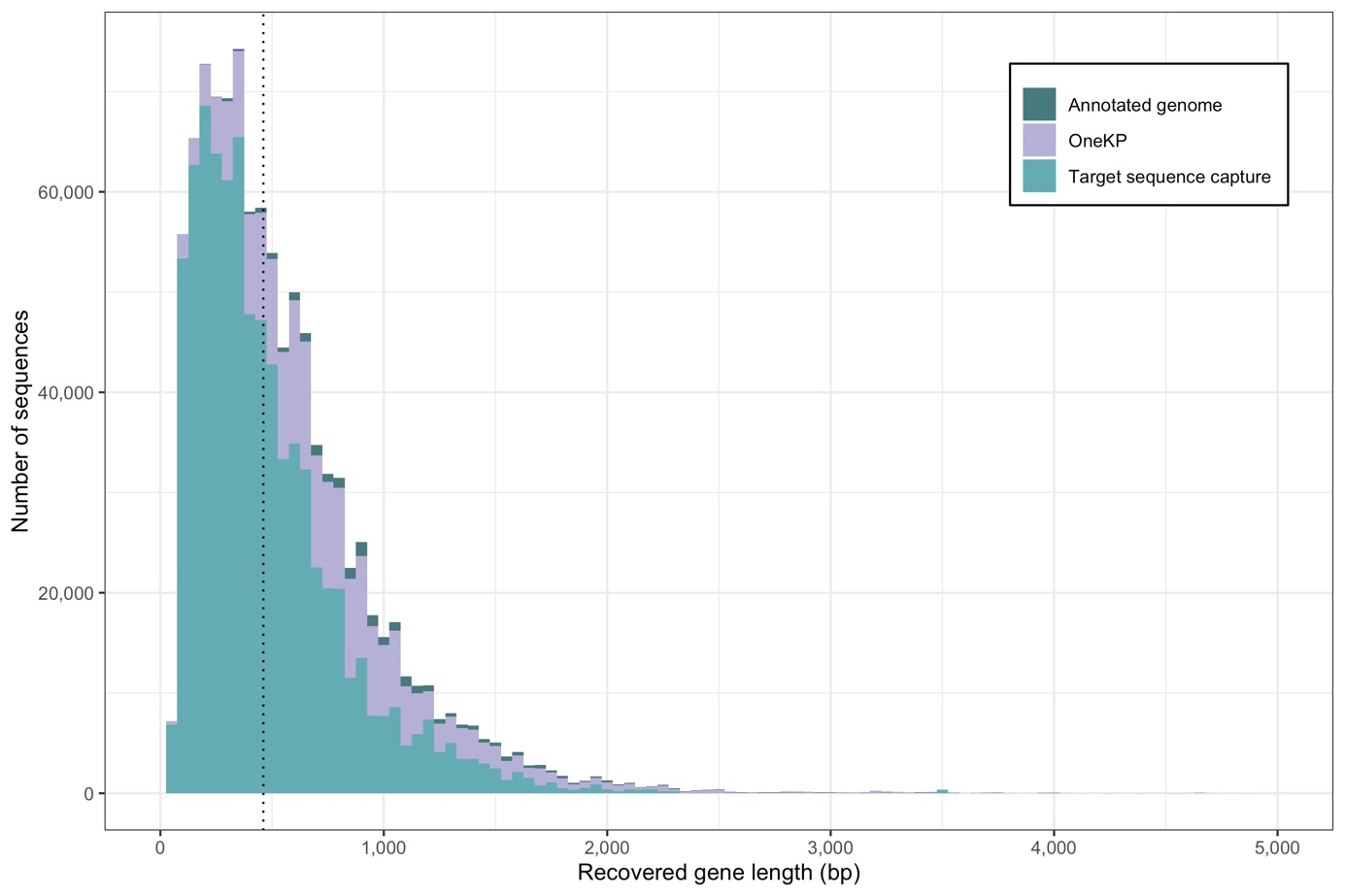

### List of Supplementary Tables

All the Supplementary Tables are available in a single file ‘Supplementary Tables S1-S5.xls’, with each Table in a separate tab.

#### **Supplementary Table S1**

Table of samples in the preliminary tree. Contains the taxonomy, validation results, recovery statistics, voucher information and ENA accession ID.

See tab ‘Table S1’.

#### **Supplementary Table S2**

Summary statistics for gene alignments and gene trees. It contains summary of alignments from AMAS and summary of gene trees statistics as node support and tree resolution.

See tab ‘Table S2’.

#### **Supplementary Table S3**

Samples from singly represented taxa. Each row represents one sample. The columns contain 1. Scientific name; 2. Name of repository where sequence is available (either ‘INSDC’ or ‘1KP’ for data stored in the International Sequence Database Collaboration and the One Thousand Plant Transcriptome Project data repository, respectively), 3. Sequence identifier in the repository, 4. Family (if applicable) of which this sample is the sole representative, 5. Order (if applicable) of which this sample is the sole representative.

See tab ‘Table S3’.

#### **Supplementary Table S4**

Samples from poorly resolved taxa. Each row represents one sample. The columns contain 1. Scientific name, 2. Name of repository where sequence is available (either ‘INSDC’ or ‘1KP’ for data stored in the International Sequence Database Collaboration and the One Thousand Plant Transcriptome Project data repository, respectively), 3. Sequence identifier in the repository, 4. Poorly resolved family (if applicable) to which this sample belongs, 5. Poorly resolved order (if applicable) to which this sample belongs.

See tab ‘Table S4’.

#### **Supplementary Table S5**

Samples which are outliers of well resolved taxa. Each row represents one sample. The columns contain 1. Scientific name, 2. Name of repository where sequence is available (either ‘INSDC’ or ‘1KP’ for data stored in the International Sequence Database Collaboration and the One Thousand Plant Transcriptome Project data repository, respectively), 3. Sequence identifier in the repository, 4. Well resolved family (if applicable) of which this sample is an outlier, 5. Well resolved order (if applicable) of which this sample is an outlier.

See tab ‘Table S5’.
