## Supplementary figures and images for "A Comprehensive Phylogenomic Platform for Exploring the Angiosperm Tree of Life"

### Supplementary Fig. S2.pdf

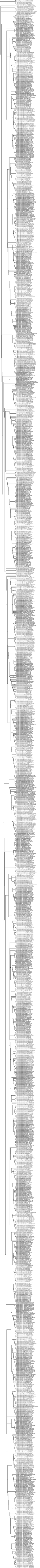
